# H3 dopaminylation and CaMKII modulate diffuse midline glioma response to CDK9 inhibition

**DOI:** 10.64898/2026.04.07.714507

**Authors:** Rebecca L. Murdaugh, Brittany R. Eberl, Rosemary U. Richard, Ena I. Campos-Hensley, Adanze N. Nnyagu, W. Austin Elam, Ai-Ni Tsao, Jack R. Tremblay, Ruiying Ma, Alfred K. Dei-Ampeh, Kieu Pham, Daniel C. Kraushaar, Kwanha Yu, Calla M. Olson, Akdes Serin-Harmanci, Benjamin Deneen, Jamie N. Anastas

## Abstract

Aberrant neurotransmitter signaling and transcriptional dysregulation are hallmarks of gliomagenesis and represent potential therapeutic targets. Monoamine neurotransmitters such as dopamine and serotonin primarily activate GPCRs but can also function epigenetically as histone H3 modifications. Here, we uncover mechanisms of crosstalk between monoamine neurotransmitter signaling, H3 dopaminylation, and RNA polymerase II (Pol2) transcription in diffuse midline glioma (DMG). We find that co-treatment with Pol2-targeting CDK9 inhibitors (CDK9i) and FDA-approved neuropsychiatric drugs, including selective serotonin reuptake inhibitors (SSRIs), synergistically reduces DMG growth. Mechanistically, CDK9i+SSRI treatment alters H3 dopaminylation patterns and represses synaptic and neurodevelopmental gene transcription associated with CDK9i resistance. Further phospho-proteomic analyses show that CDK9i monotherapy activates pro-survival CaMKII signaling, which can be suppressed by co-treatment with neuromodulatory drugs. These studies establish roles for H3 dopaminylation and neurotransmitter signaling in DMG gene regulation and response to CDK9i, suggesting that monoamine neurotransmitter pathways may be exploited as a therapeutic strategy for DMG.

## Main

Diffuse midline glioma (DMG) is an aggressive subtype of pediatric high-grade glioma with a five-year survival rate of <5%^1^. More than 80% of DMG tumors exhibit H3K27M driver mutations^2^, which induce abnormal RNA polymerase II (Pol2)-dependent transcription that may sensitize DMG to Pol2-targeting drugs^3^. In addition to transcriptional dysregulation, aberrant neurotransmitter signaling is emerging as another potential therapeutic target in glioma^4^.

Monoamine neurotransmitters like dopamine and serotonin signal through G protein-coupled receptors (GPCRs) to regulate downstream kinases, including PKA and CaMKII, which are often dysregulated in cancer^5,6^. Previous studies find that altered neurotransmitter signaling regulates glioma membrane depolarization and downstream kinase signaling to stimulate or inhibit tumorigenesis in a context-dependent manner^7–12^. Monoamine neurotransmitters can also be covalently attached to proteins like histone H3^13–15^ and can therefore affect both aberrant GPCR signaling and epigenetic dysregulation in cancer. While altered histone monoaminylation is known to regulate gene expression and tumorigenesis in pancreatic cancer, colorectal cancer, and ependymoma^16,17^, the potential impact of histone monoaminylation on DMG chromatin regulation, transcription, and response to therapy remains unexplored.

Cyclin-dependent kinase 9 (CDK9) regulates transcription by phosphorylating serine 2 of the C-terminal domain (CTD) repeats of the largest subunit of Pol2 (pS2-Pol2) to initiate Pol2 elongation^18,19^. CDK9 inhibitors (CDK9i) have demonstrated efficacy in pre-clinical models of DMG^3^, and early-stage clinical trials using the CDK9i zotiraciclib (ZTR) for the treatment of adult brain tumors are underway (NCT02942264, NCT05588141), incentivizing the further optimization of CDK9-targeting strategies for brain tumor treatment^20^. However, DMG cells exhibit both intrinsic and acquired resistance to targeted therapies^21^, and dose-limiting toxicities of CDK9 inhibitors may limit their therapeutic potential as monotherapies^22^.

We describe a unique pattern of H3 dopaminylation at H3K27M-occupied genes encoding Pol2-transcription regulators, suggesting a role for neurotransmitter-chromatin signaling in DMG gene regulation. Additional studies identify neurotransmitter signaling pathways as markers of CDK9i resistance and demonstrate that co-targeting Pol2 transcription with CDK9i and neurotransmitter signaling with FDA-approved neuropsychiatric drugs synergistically reduces DMG growth *in vitro* and reduces tumor growth in xenograft models. Subsequent phospho-proteomic, transcriptomic, and epigenomic profiling studies suggest that these neurotransmitter-targeting drugs synergize with CDK9i in DMG cells through the modulation of CaMKII signaling and histone dopaminylation. Together, these findings establish neurotransmitter-chromatin signaling as a key regulator of Pol2 transcriptional programs and therapeutic responses in DMG.

## Results

### H3 dopaminylation marks genes encoding Pol2 transcriptional regulators in DMG

Histone H3 monoaminylation by serotonin and dopamine (H3Q5ser, H3Q5dop) is associated with aberrant transcription in ependymoma and other cancers^16,23,24^, but a role for histone monoaminylation in DMG gene regulation is still undescribed. To explore neurotransmitter to chromatin signaling in DMG, we first performed targeted mass spectrometry (MS) on four DMG cell lines, revealing that DMG cells produce multiple neurotransmitters, including dopamine, histamine, and serotonin (Fig. **1a**). As dopamine was the most abundant monoamine neurotransmitter in the MS analysis, we asked whether histones undergo dopaminylation in DMG. Immunofluorescence staining of DMG cell lines using an antibody recognizing H3 co-modified by H3K4me3 and Q5 dopaminylation (H3K4me3Q5dop) confirms the presence of H3 dopaminylation in multiple DMG cell lines (Fig. **1b**).

**Fig. 1:**
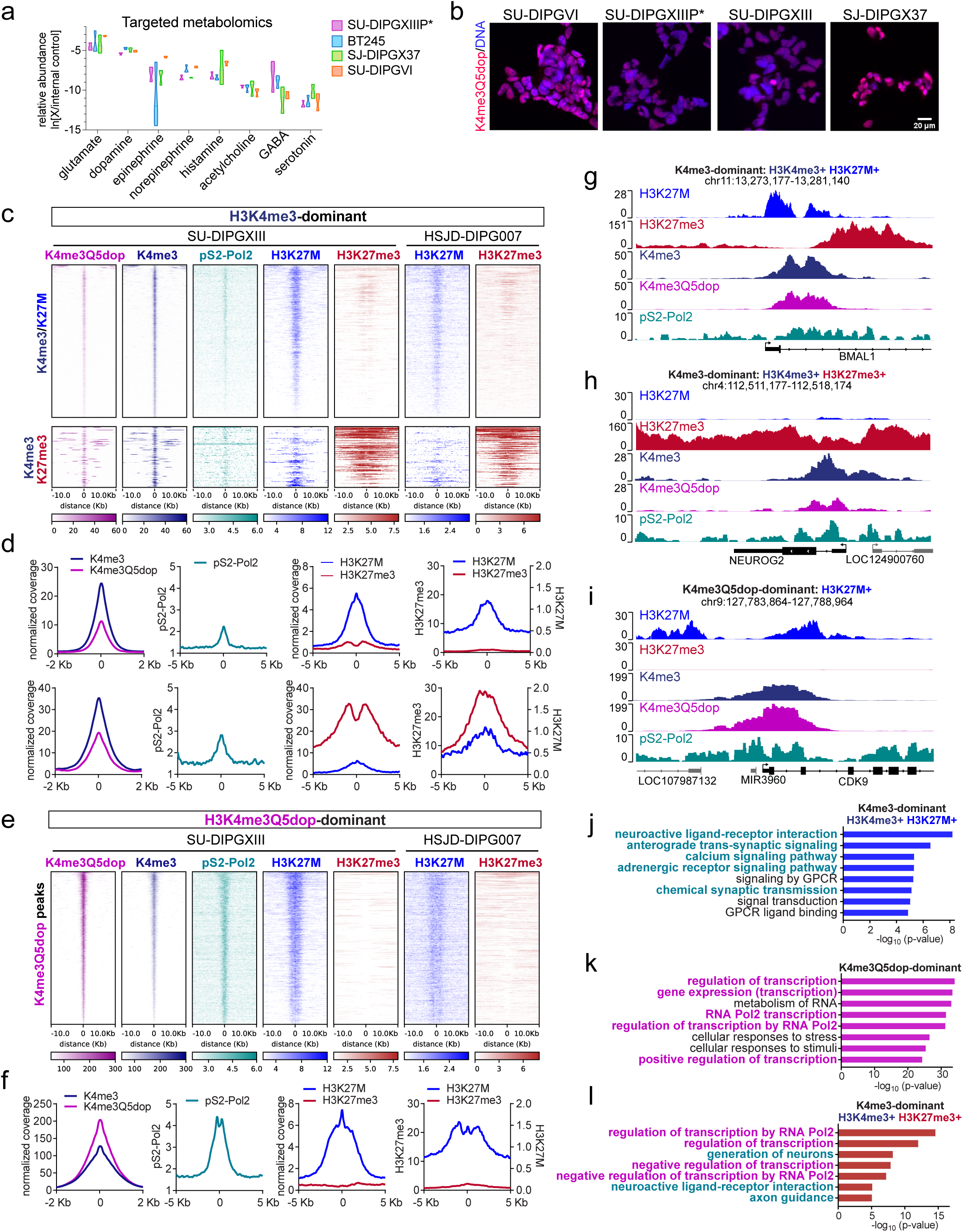
H3 dopaminylation marks genes encoding Pol2 transcriptional regulators in DMG. a, Relative abundance of neurotransmitters in 4 DMG cell lines (SU-DIPGXIIIP*, BT245, SJ-DIPGX37, SU-DIPGVI) determined by targeted metabolomics. For **a**, *n* = 3 replicates per cell line. **b**, Representative images of H3K27M mutant DMG cells stained to detect H3K4me3Q5dop. Scale bar is 20 μm. **c**-**f**, Heatmaps (**c**,**e**) and average profile plots (**d**,**f**) from CUT&RUN analysis of H3K4me3, H3K4me3Q5dop, and pS2-Pol2 in SU-DIPGXIII aligned to previously published H3K27M and H3K27me3 profiling data from SU-DIPGXIII and HSJD-DIPG007 cells. Panels **c**,**d** show H3K4me3-dominant peaks that co-localized either with H3K27M (top panels) or with H3K27me3 (bottom panels) and panels **e**,**f** show H3K4me3Q5dop-dominant peaks co-occupied by H3K27M. **g*-*i**, Genome browser snapshots showing low H3 dopaminylation and H3K4me3/H3K27M enrichment at the *BMAL1* gene (**g**), low H3 dopaminylation and a bivalent chromatin signature (H3K4me3/K27me3) at the *NEUROG2* locus (**h**), and strong H3K4me3Q5dop and H3K27M co-enrichment at the *CDK9* gene locus (**i**). **j*-*l**, Enriched gene categories from KEGG, REACTOME, and GO (biological process) gene sets databases among the H3K27M+/H3K4me3-dominant peaks (**j**), the H3K27M+/H3K4me3Q5dop-dominant peaks (**k**), or the H3K27me3/H3K4me3 bivalent chromatin peaks (**l**). Gene sets related to Pol2 transcriptional regulation are shown in teal and related to neuronal signaling are shown in purple.

To profile histone dopaminylation genome-wide, we then performed CUT&RUN analysis on SU-DIPGXIII cells, revealing co-localization of H3K4me3Q5dop, H3K4me3, and elongating RNA polymerase II (Pol2) marked by CTD serine-2 phosphorylation (pS2-Pol2) (Extended Data Fig. **1a**). Further analyses identified genomic regions marked by either a stronger H3K4me3 signal compared to H3K4me3Q5dop (H3K4me3-dominant; Fig. **1c,d**) or associated with increased H3K4me3Q5dop compared to H3K4me3 binding (H3K4me3Q5dop-dominant, Fig. **1e,f**). Consistent with previous studies suggesting a permissive role for histone monoaminylation in Pol2 transcription^15,25^, we observe increased pS2-Pol2 occupancy at the H3K4me3Q5dop-dominant peaks (Fig. **1c-f**).

The DMG chromatin landscape is characterized by H3K27M oncohistone binding associated with gene activation and largely non-overlapping regions of H3K27me3 at tumor suppressor genes that are retained despite H3K27M-dependent inhibition of PRC2 activity^26–28^. To determine the relationship between H3 dopaminylation and H3K27M and H3K27me3 chromatin binding patterns in DMG, we aligned these H3K4me3-and H3K4me3Q5dop-dominant peaks with published H3K27M and H3K27me3 chromatin profiles from SU-DIPGXIII and HSJD-DIPG007 cells^29^, revealing specific patterns of H3K27M and H3K27me3 incorporation at the H3K4me3-and H3K4me3Q5dop-dominant loci (Supplementary Table 1). Most of the H3K4me3-dominant peaks exhibited co-enrichment with H3K27M oncohistones, as seen at the *BMAL1* gene, which encodes a transcription factor linked to circadian rhythms^30^ (2,264 regions, top panels; Fig. **1c,d** and **g**). A smaller subset of the H3K4me3-dominant peaks showed co-enrichment with H3K27me3 and low H3K27M binding, suggestive of a bivalent H3K4me3/H3K27me3 chromatin state linked to the regulation of progenitor cell and differentiation genes in DMG and other cancers^26,31^ (566 regions, bottom panels; Fig. **1c,d**). Representative loci exhibiting this pattern of low H3K4me3Q5dop enrichment and H3K4me3/H3K27me3 bivalency include genes encoding calcium regulators (*CALCA* and *CALCR*; Extended Data Fig. **1b,c**) and Neurogenin-2 (*NEUROG2*) (Fig. **1h**), which encodes a pro-neurogenic transcription factor^32^.

In contrast, the H3K4me3Q5dop-dominant peaks co-localized with H3K27M and H3K4me3 but showed minimal H3K27me3 enrichment, suggesting a correlation between histone dopaminylation and an active chromatin state in DMG (2,660 regions; Fig. **1e,f**). This pattern of H3K4me3Q5dop/H3K27M co-enrichment was observed at genes encoding Pol2 transcription regulators, including cyclin-dependent kinases 9 and 12 (*CDK9* and *CDK12*; Fig. **1i** and Extended Data Fig. **1d**), which regulate Pol2 pause release and elongation dynamics^33^ and at additional Pol2 co-activator genes like *UBTF*^34^ and *TADA1*^35^ (Extended Data Fig. **1e,f**). Consistently, gene ontology (GO) analysis reveals that the H3K4me3-dominant/H3K27M+ peaks are enriched near genes associated with calcium signaling, neuroactive ligand receptor interactions, and synaptic transmission (Fig. **1j**), whereas the H3K4me3Q5dop-dominant peaks are found near genes involved in Pol2 transcription (Fig. **1k**). Finally, the H3K4me3-dominant, bivalent genes encode both Pol2 regulators and neuronal signaling and axon guidance factors (Fig. **1l**). Together, these analyses reveal distinct patterns of histone dopaminylation, H3K4me3, H3K27me3, and H3K27M incorporation at genes encoding Pol2 and neuronal signaling modulators, suggesting potential roles for histone monoaminylation in DMG transcriptional dysregulation and response to neurotransmitters.

### Neuronal signaling genes are upregulated in CDK9i-resistant DMG xenograft tumors

Our finding that H3 dopaminylation marks *CDK12*, *CDK9*, and other Pol2 regulatory genes points to a potential connection between neurotransmitter signaling and transcriptional regulation in DMG. Supporting this idea, we find that neuronal signaling genes are upregulated in DMG xenograft tumors that persisted following chronic treatment with zotiraciclib (ZTR), a Pol2-targeting CDK9 inhibitor (CDK9i) under investigation as a treatment for adult glioma (NCT05588141). Although ZTR selectively kills H3K27M mutant DMG cells *in vitro* (Extended Data Fig. **2a-c**), 3 weeks of treatment with a sub-threshold dose of ZTR (15 mg/kg, p.o.) only transiently reduced the growth of pontine SU-DIPGXIIIP*+ZsGreen/luciferase xenograft tumors without extending overall survival (Fig. **2a-c**), suggesting that CDK9i resistance develops *in vivo*. RNA-seq analysis reveals increased expression of transcripts involved in glutamate signaling, potassium channel activity, and neuronal projections in the ZTR-resistant versus vehicle-treated tumors (Fig. **2d,e**, Extended Data Fig. **2d**, and Supplementary Table 2), indicating that altered neuronal signaling may contribute to CDK9i resistance.

**Fig. 2:**
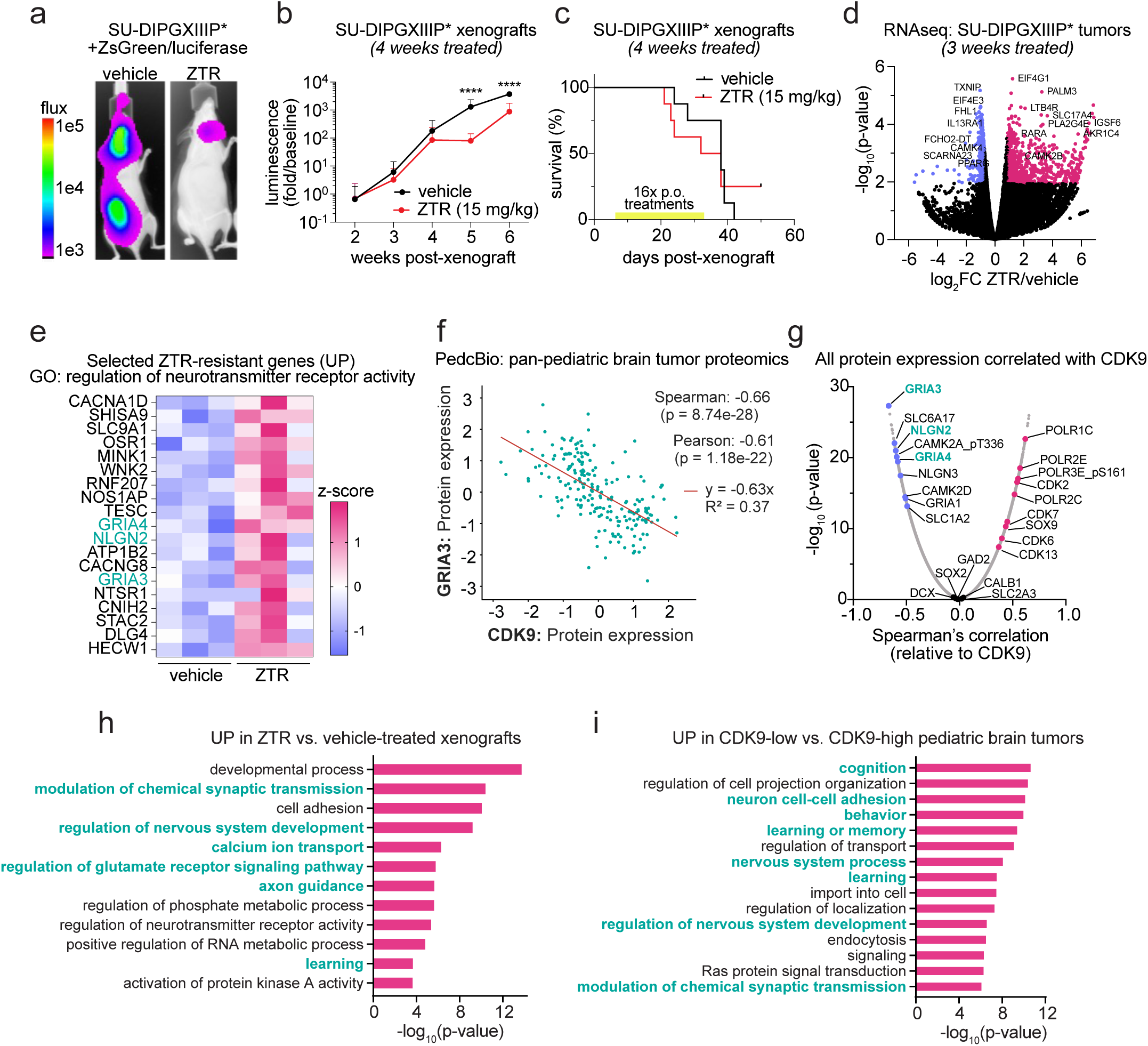
Neuronal signaling genes are upregulated in CDK9i-resistant DMG xenograft tumors. **a**,**b**, Representative images (**a**) and quantification (**b**) of tumor luciferase signal (BLI) from orthotopic xenografts of SU-DIPGXIIIP*+ZsGreen/luciferase cells following treatment with vehicle or the ZTR (15 mg/kg PO, 4 days/week). **c**, Kaplan Meier survival analysis comparing vehicle-and ZTR-treated mice bearing SU-DIPGXIIIP* xenografts for 4 weeks (yellow bar shows treatment period). For **b**,**c**, *n* = 8 mice per group. **d**, Volcano plot summarizing bulk RNA-seq analysis of SU-DIPGXIIIP* xenograft cells isolated from dissociated mouse brains after three weeks of treatment with vehicle or ZTR. In panel **d**, *n* = 3 mice per group. **e**, Heatmap of RNA-seq Z-scores of neurotransmitter receptor signaling genes that are differentially expressed in ZTR-resistant DMG tumors. **f**, Scatter plot showing a negative correlation between GRIA3 and CDK9 protein expression in a pan-pediatric brain tumor proteomics dataset (PedcBio, CPTAC) with a linear regression line shown in red. **g**, Volcano plot showing the ranked Spearman’s correlation coefficients of all proteins relative to CDK9 protein expression (x-axis) and the-log_10_(*p*-value) from the pediatric glioma proteomics data. **h,i,** Enriched gene ontology (GO) terms associated with transcripts upregulated at the mRNA level in ZTR-treated SU-DIPGXIIIP* xenografts (**h**) or at the protein level in CDK9-low pediatric brain tumor samples (**i**).

A complementary analysis of pediatric glioma patient tumor proteomics data^36^ accordingly reveals a negative association between CDK9 protein abundance and the expression of neuronal signaling proteins (Supplementary Table 2). For example, glutamate receptors 3 and 4 (*GRIA3 and GRIA4*) and neuroligin 2 (*NLGN2*) were up-regulated in both ZTR-treated xenograft tumors (Fig. **2e**) and in CDK9-low patient samples (Fig. **2f,g**). GO analysis across both datasets further confirms that genes associated with neuronal signaling and brain development are upregulated at a transcriptional level in the ZTR-resistant DMG xenograft tumors (Fig. **2h**) and similarly elevated at a protein level in CDK9-low versus CDK9-high patient tumors (Fig. **2i**). These results suggest that neuronal signaling pathways are dysregulated by chronic CDK9 inhibition or CDK9 downregulation, suggesting crosstalk between neurotransmitter signaling and transcriptional dysregulation in DMG.

### FDA-approved drugs targeting neurotransmitter signaling sensitize DMG to CDK9i

Additional evidence for a functional relationship between aberrant neurotransmitter signaling and Pol2 regulation in DMG comes from a drug synergy screen that identified neurotransmitter-targeting drugs as potent CDK9i-synergizers. We screened a library of FDA-approved and bioactive compounds on top of an IC_25_ dose of ZTR and monitored DMG cell growth by CellTiter-Glo. We find that 55% (12/22) of the compounds that enhanced SU-DIPGXIII response to low-dose ZTR treatment modulate neuronal signaling pathways (Fig. **3a,b**, Extended Data Fig. **3a**, and Supplementary Table 3). ZTR-synergistic neurotransmitter-targeting drugs identified in this screen include ion channel blockers (bepridil, mefloquine), neurotransmitter GPCR antagonists (tolterodine, clemastine, asenapine, aprepitant, silodosin, prochlorperazine, nebivolol, meclizine), and selective neurotransmitter reuptake inhibitors (sertraline, duloxetine) (Fig. **3a,b**). None of these synergistic drugs significantly reduced DMG growth on their own, even at the highest dose (5 µM), yet treatment with these compounds enhanced SU-DIPGXIII sensitivity to ZTR.

**Fig. 3:**
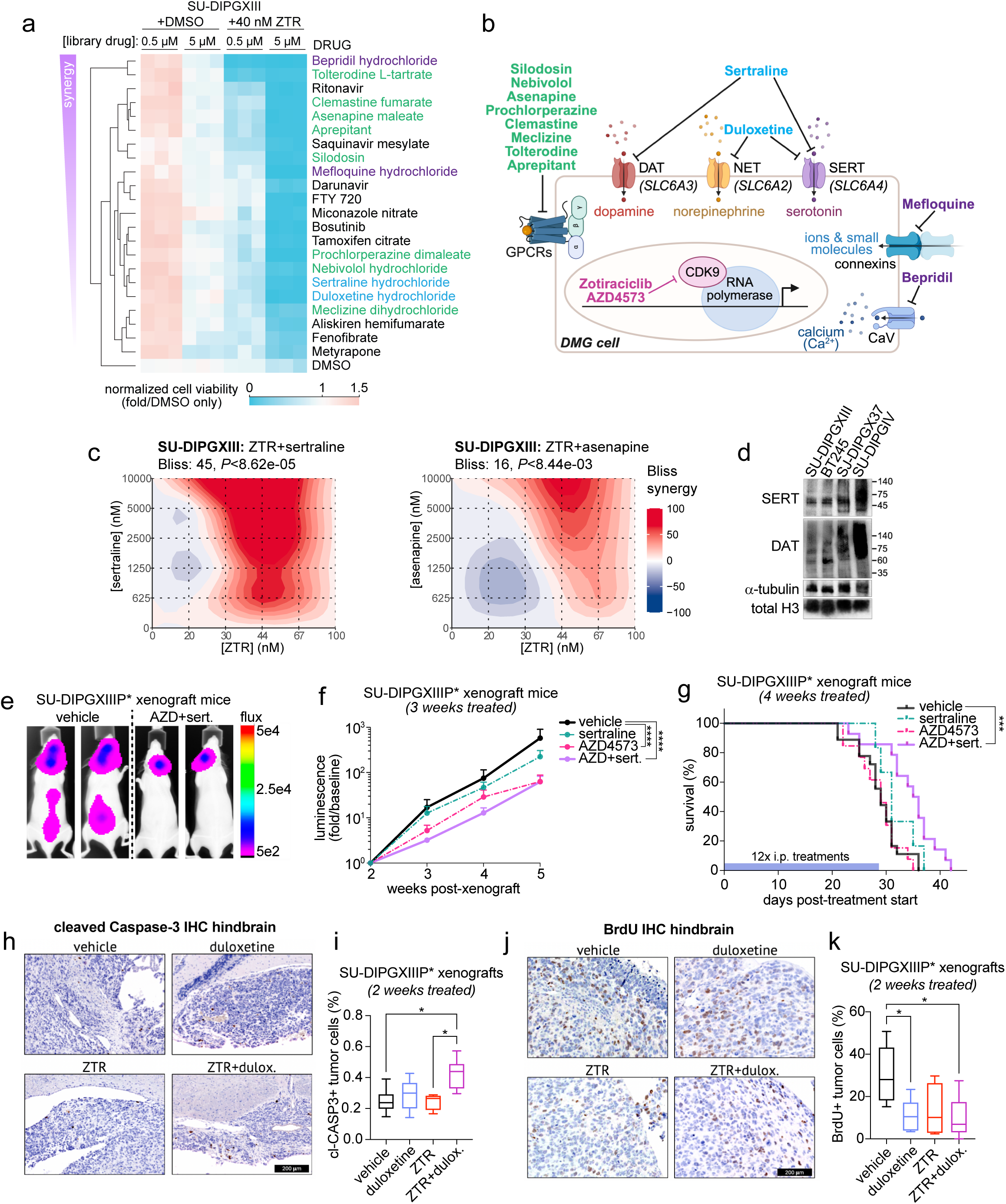
FDA-approved drugs targeting neurotransmitter signaling sensitize DMG to CDK9i. a, Fold change in SU-DIPGXIII cell viability (CellTiter-Glo) after 7 days treatment with a 0.5 µM and 5 µM dose of an FDA-approved drug library combined with either vehicle (DMSO) or 40 nM ZTR (IC_25_ dose) for ZTR-synergistic hits from drug screen. **b,** Diagram depicting CDK9i-synergistic FDA approved drugs and their molecular targets related to neuronal signaling (with ion channel blockers shown in purple, neurotransmitter GPCR antagonists shown in green, and SSRI/SNRI shown in blue. **c**, Bliss synergy scores derived from SU-DIPGXIIII cells treated with dose curve matrices of ZTR combined with either the SSRI, sertraline (left panel), or the neurotransmitter GPCR antagonist, asenapine (right panel). **d**, Western blots validating the expression of neurotransmitter reuptake transporters in DMG cell lines. **e,f**, Representative tumor BLI signals (**e**) and the average fold change in BLI signal (**f**) from SU-DIPGXIIIP* xenografts treated with the CDK9i AZD4573 (2.5 mg/kg) and/or sertraline (2.5 mg/kg) 3 times per week for 3 weeks. **g**, Kaplan-Meier survival analysis indicating that treatment with a combination of AZD4573 and sertraline significantly increased survival in the SU-DIPGXIIIP* xenograft model (blue bar shows treatment period; *P*<0.0008, Mantel-Cox log-rank test). For **f**,**g**, *n* = 18 mice for vehicle group, *n* = 6 mice for sertraline group, *n* = 13 mice for AZD4573 group, and *n* = 14 mice for AZD4573+sertraline group (AZD+sert.). **h-k**, Representative images and quantification of cleaved Caspase-3 (cl-CASP3) positive (**h,j**) or BrdU positive (**i**,**k**) SU-DIPGXIIIP* xenograft tumor cells following 2 weeks treatment with duloxetine (5 mg/kg) and ZTR (20 mg/kg) dosed 3 days per week. For **h**-**k**, *n* = 6 mice for vehicle group, *n* = 5 mice for duloxetine group, *n* = 4 mice for zotiraciclib (ZTR) group, and *n* = 6 mice for the zotiraciclib+duloxetine group (ZTR+dulox.).

To validate the synergistic activity of CDK9i combined with neurotransmitter-targeting drugs, we set up dose curve matrices in SU-DIPGXIII, SU-DIPGIV, and SJ-DIPGX37 cells. We observe a synergistic effect (Bliss score >10)^37^ following co-treatment with ZTR and sertraline, an SSRI^38^, with duloxetine, an SNRI, or with asenapine, an antagonist of GPCRs for serotonin, dopamine, and other neurotransmitters^39^ (Fig. **3c** and Extended Data Fig. **3b**). Further validation studies show that multiple DMG cell lines (SU-DIPGXIII, BT245, SJ-DIPGX37) are sensitive to combinations of structurally distinct CDK9i (ZTR or AZD4573) and several SSRI/SNRI (sertraline, duloxetine, or escitalopram) as determined through manual cell counts or CellTiter-Glo assays (Extended Data Fig. **3c,d**). Consistent with on-target activity, immunoblotting and immunofluorescence staining analyses verified the expression of the known neurotransmitter reuptake transporters (SERT, DAT, NET) targeted by these SSRI/SNRIs (Fig. **3d** and Extended Data Fig. **3e**).

### Co-targeting neurotransmitter signaling and CDK9 reduces DMG xenograft growth

We next asked whether CDK9i and SSRI/SNRI drug combinations can disrupt DMG xenograft progression. To assess on-target activity of these drugs *in vivo*, we collected brains from mice bearing pontine SU-DIPGXIIIP* tumors following 4 weeks of treatment with low doses of the CDK9i AZD4573 (2.5 mg/kg), sertraline (2.5 mg/kg), or a combination (3 days/week, IP) and performed staining for pS2-Pol2, a known CDK9 substrate^40,41^. Treatment with AZD4573 alone, sertraline alone, and AZD4573+sertraline combination all significantly reduced pS2-Pol2 staining in tumor tissue but did not affect pS2-Pol2 staining intensity in adjacent normal brain tissue (Extended Data Fig. **3f,g**). Long-term treatment studies reveal that, although both AZD4573 alone and AZD4573+sertraline combination treatment reduced DMG tumor volume by bioluminescent imaging (BLI; Fig. **3e,f**), only the AZD4573+sertraline combination therapy was sufficient to increase survival (Fig. **3g**).

To further characterize the effect of CDK9i combination therapy on DMG cell proliferation and death *in vivo*, we BrdU-pulsed SU-DIPGXIIIP* xenograft mice and performed IHC on brains collected following treatment with vehicle, the SNRI duloxetine (5 mg/kg), ZTR (20 mg/kg), or ZTR+duloxetine dosed 3 days per week for 2 weeks. We observe a higher frequency of cleaved Caspase-3+ (cl-CASP3) dead or dying tumor cells in mice treated with ZTR+duloxetine relative to the vehicle and ZTR only conditions (Fig. **3h,i**) and a reduction in BrdU+ S-phase cells in the duloxetine alone and ZTR+duloxetine combination-treated tumors relative to vehicle controls (Fig. **3j,k**). These data showing that DMG are vulnerable to combinations of drugs targeting neurotransmitter signaling and CDK9 *in vivo* provide additional evidence that monoamine neurotransmitter signaling modulates DMG response to CDK9i-induced Pol2 transcriptional stress.

### CDK9i+SSRI co-treatment silences oncogenic and neuronal signaling genes enriched in CDK9i-resistant DMG

To better understand mechanisms of CDK9i+SSRI drug synergy in DMG, we performed bulk RNA-seq on SU-DIPGXIII cells treated with CDK9i and sertraline either alone or in combination (Supplementary Table 4). Principal component analysis (PCA) of our RNA-seq data shows clustering of experimental replicates and a greater difference in both PC1 and PC2 of the ZTR+sertraline-treated samples compared to samples treated with single drugs or vehicle (Fig. **4a**). Accordingly, ZTR+sertraline combination treatment leads to a greater number of differentially expressed genes (DEGs; log2FC+/-1.5, adjusted *P*<0.01) than either ZTR or sertraline alone (Extended Data Fig. **4a**). Co-treatment with sertraline and a second CDK9i (AZD4573+sertraline) similarly induces more robust transcriptional changes than AZD4573 alone (Fig. **4a**, Supplementary Table 4 and Extended Data Fig. **4a**) with a high degree of overlap between the DEGs identified in the ZTR+sertraline and AZD4573+sertraline treatment groups (Fig. **4b**), demonstrating that two structurally distinct CDK9i produce similar transcriptional outputs with SSRI co-treatment.

**Fig. 4:**
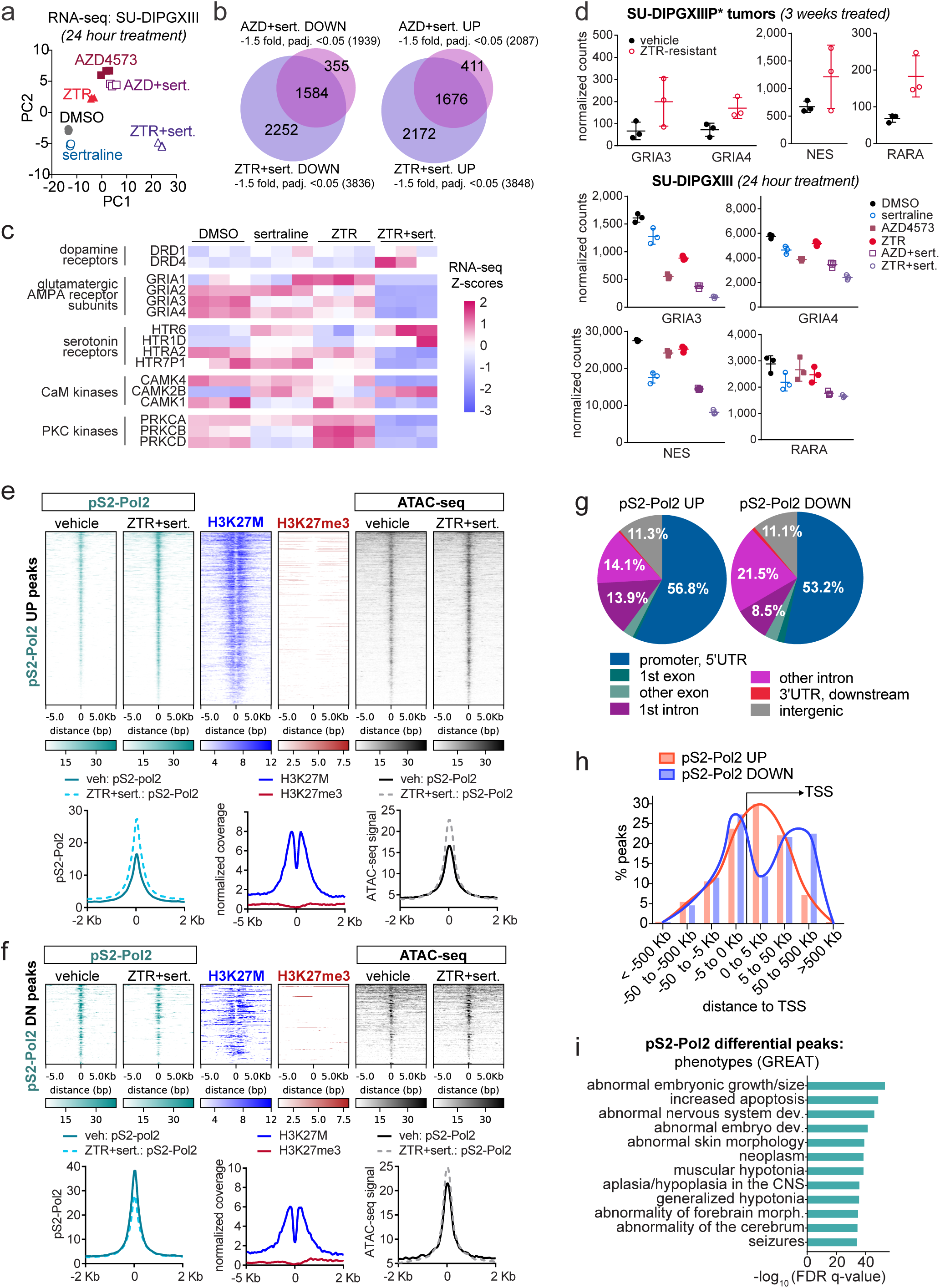
CDK9i+SSRI co-treatment silences oncogenic and neuronal signaling genes enriched in CDK9i-resistant DMG. **a**, PCA analysis of RNA-seq data from SU-DIPGXIII cells treated with vehicle, 20 nM ZTR, 6 nM AZD4573 (AZD), 5 µM sertraline (sert.), ZTR+sert., or AZD+sert. for 24 hours. **b**, Venn diagrams illustrating the overlap between downregulated (left) and upregulated genes (right) after treatment with ZTR+sert. or with AZD4573+sert. (+/-1.5-fold, adjusted *P*<0.05). **c**, Heatmap of Z-scores for transcripts that were uniquely upregulated or downregulated by ZTR+sert. combination treatment highlighting differential expression of glutamate, dopamine, and serotonin receptors. **d**, Normalized transcript counts for selected differentially expressed genes (*GRIA3*, *GRIA4*, *RARA*, *NES*) that were upregulated in the ZTR-resistant DMG tumor RNA-seq dataset (see also Fig. 2d**,e**; Supplementary Table 1), but conversely downregulated in SU-DIPGXIII cells co-treated with CDK9i+sertraline for 24 hours. **e,f**, Heatmaps from CUT&RUN profiling in SU-DIPGXIII cells treated for 24 hours with either vehicle (DMSO) or ZTR+sert showing either increased (**e**) or decreased (**f**) pS2-Pol2 binding due to ZTR+sert. treatment. In panels **e,f,** differential pS2-Pol2 co-localize with H3K27M and exhibit differential chromatin accessibility by ATAC-seq (see also Extended Data Fig. 5 and Supplementary Table 5). **g,h**, Frequency of pS2-Pol2-up or-down (DN) peaks within annotated genomic elements (**g**) and their relative distances to the nearest transcription start site (TSS) (**h**). **i**, Significant GO terms related to central nervous system development and embryonic phenotypes enriched among genes located nearest to the differential pS2-Pol2 peaks (combining increased & decreased peaks).

Further GO analyses reveal that sertraline treatment upregulates transcripts involved in extracellular matrix organization, whereas CDK9i treatment induces genes involved in dopamine metabolism and axon guidance (Extended Data Fig. **4b,c**). Combined ZTR+sertraline treatment induces pronounced changes in the expression of genes involved in neurodevelopment and synaptic signaling (Extended Data Fig. **4d**). These ZTR+sertraline-induced transcriptional changes include repression of PKC subunit and glutamate receptor transcripts (*PRKCA*, *PRKCB*, *PRKCD,* and *GRIA1*-*4*) and altered expression of dopamine and serotonin receptors (Fig. **4c**). Finally, comparing these RNA-seq results to our previous analysis of ZTR-resistant xenograft tumors reveals that a subset of transcripts induced by chronic ZTR treatment *in vivo* (Fig. **2d,e** and Supplementary Table 2) were inhibited by CDK9i+SSRI co-treatment, including glutamate receptors (*GRIA3*, *GRIA4*), *Nestin* (*NES)*, and the *Retinoic Acid Receptor Alpha* (*RARA)* mRNAs (Fig. **4d**). These analyses suggest that CDK9i+SSRI co-treatment synergistically alters the expression of neuronal signaling genes in DMG, including transcripts linked to CDK9i resistance.

To define mechanisms underlying altered Pol2 transcriptional regulation induced by CDK9-and neurotransmitter-targeted combination therapy, we performed CUT&RUN for pS2-Pol2 and ATAC-seq after 24 hours of treatment with ZTR+sertraline or vehicle control. CDK9i+SSRI combination treatment yielded 2,001 upregulated and 869 downregulated pS2-Pol2 peaks (*P*<0.05; Fig. **4e,f** and Supplementary Table 5) as well as 2,458 upregulated and 2,215 downregulated ATAC-seq peaks (*P*<0.05, log2FC+/-1; Extended Data Fig. **5a,b** and Supplementary Table 5). Integrating our ATAC-seq and RNA-seq results shows that ZTR+sertraline treatment modifies chromatin accessibility at 341 upregulated genes and at 443 downregulated genes (Extended Data Fig. **5c**), suggesting that CDK9i+SSRI causes localized remodeling of chromatin structure and disrupts Pol2 regulation at downstream targets.

Additional analyses reveal that both the ZTR+sertraline-induced and-repressed pS2-Pol2 peaks co-localized with H3K27M oncohistones (Fig. **4e,f**) and were enriched at promoters and gene bodies (Fig. **4g**). A closer examination of the pS2-Pol2 chromatin profiles reveals a reduced distance between upregulated pS2-Pol2 peaks and the nearest transcription start sites (TSS) compared to the pS2-Pol2-down peaks (Fig. **4h**). These results suggest that CDK9i+SSRI treatment leads to pS2-Pol2 accumulation near the 5’ end of genes, consistent with reduced transcriptional elongation. Finally, GO analysis of the genes associated with differential pS2-Pol2 occupancy shows that ZTR+sertraline treatment alters Pol2 dynamics at genes related to embryogenesis, apoptosis, nervous system development, and cancer (Fig. **4i**). Together, these analyses reveal that co-targeting CDK9 and monoamine neurotransmitter signaling disrupts pS2-Pol2 localization and chromatin accessibility at H3K27M-bound genes involved in brain development and malignancy.

### CDK9i+SSRI treatment alters H3 dopaminylation at neuronal signaling and brain development genes

We next determined the consequences of CDK9i+SSRI co-treatment on the H3 dopaminylation landscape in DMG. CUT&RUN analysis identified 1,406 upregulated and 1,585 downregulated H3K4me3Q5dop peaks in SU-DIPGXIII cells treated with ZTR+sertraline compared to vehicle (*P*<0.05; Fig. **5a,b** and Supplementary Table 5). However, H3K4me3 enrichment was not significantly altered at most of the differential H3K4me3Q5dop peaks. Further analyses reveal that ZTR+sertraline increased H3K4me3Q5dop at sites with a higher ratio of H3K27me3 to H3K27M binding (Fig. **5a**), reminiscent of the H3K4me3/H3K27me3+ bivalent peaks observed in untreated SU-DIPGXIII cells (bottom panels Fig. **1c,d**), as demonstrated by a gain of H3K4me3Q5dop at bivalent genes like *NEUROG2*, *CALCA*, and *DLGAP3* (Fig. **5c** and Extended Data Fig. **5d,e**). In contrast, ZTR+sertraline treatment slightly reduced both H3K4me3Q5dop and H3K4me3 at H3K27M-enriched target genes (Fig. **5b**), including at the *GRIA4*, *MMACHC*, and *GPR19* gene loci (Fig. **5d** and Extended Data Fig. **5f,g**). Highlighting *GRIA4* as an example, we observe increased pS2-Pol2 near the TSS, indicative of Pol2 pausing, and decreased H3K4me3Q5dop enrichment (Fig. **5d**) as well as a corresponding decrease in *GRIA4* gene expression upon ZTR+sertraline treatment (Fig. **4c,d**).

**Fig. 5:**
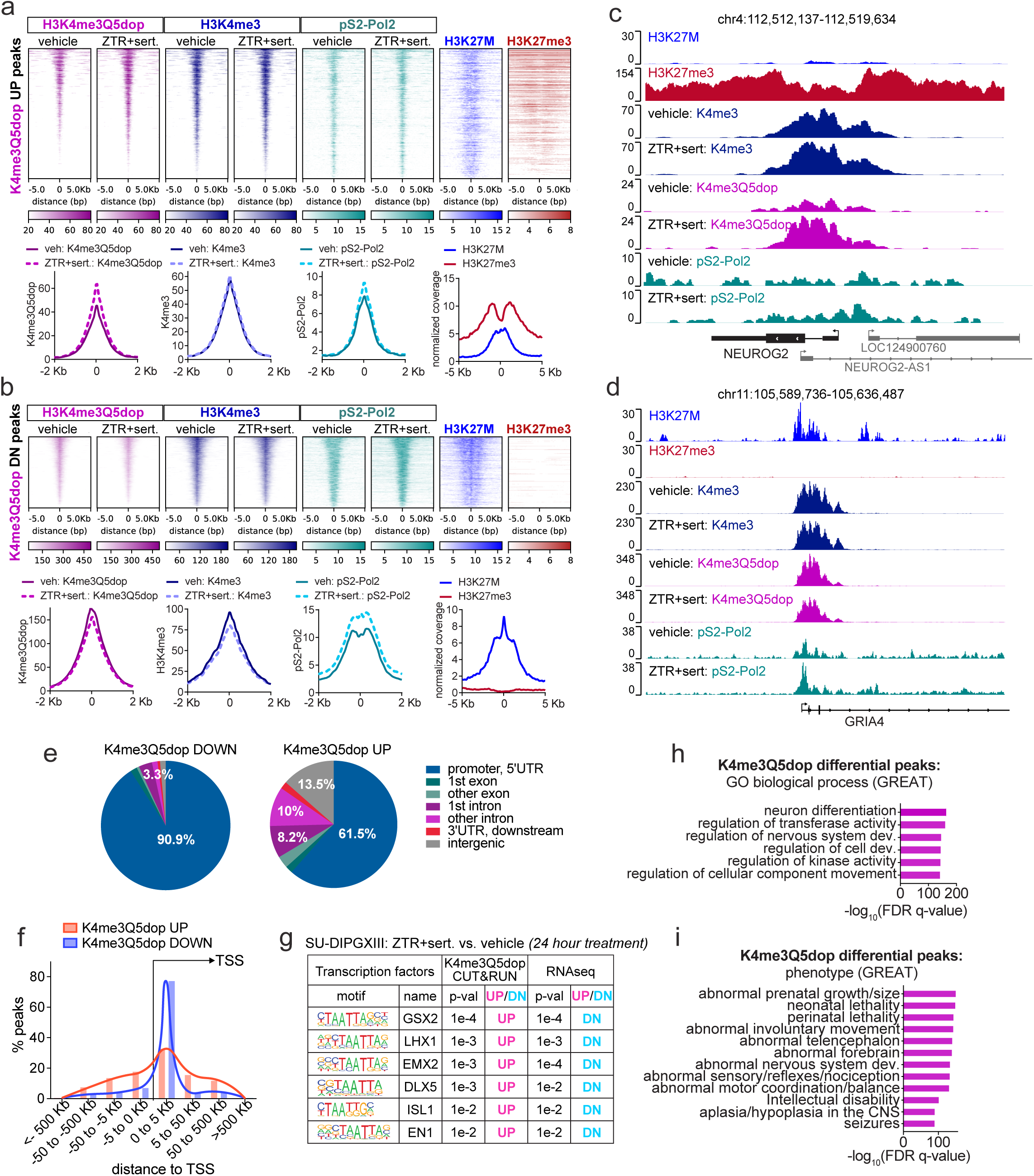
CDK9i+SSRI treatment alters H3 dopaminylation at neuronal signaling and brain development genes. **a**,**b**, Heatmaps centered around ZTR+sertraline treatment-induced (**a**) or-repressed (**b**) H3K4me3Q5dop CUT&RUN peaks aligned to H3K4me3, pS2-Pol2, H3K27M, and H3K27me3 chromatin profiling data. **c**,**d**, Genome tracks summarizing CUT&RUN chromatin profiling data at the *NEUROG2* (**c**) and *GRIA4* (**d**) genes. **e**, Frequency of H3K4me3Q5dop-up or-down (DN) peaks at different distances from the nearest TSS. **f**, Motifs enriched in the promoters of genes with decreased expression (DN) by RNA-seq and within increased H3K4me3Q5dop-up peaks in SU-DIPGXIII cells treated with ZTR+sert. versus vehicle. **g**,**h**, Significant GO terms related to biological processes (**g**) and phenotypes (**h**) for genes closest to differential H3K4me3Q5dop peaks combining both up and down peaks.

Additional analyses suggest that ZTR+sertraline treatment predominantly represses H3K4me3Q5dop enrichment near TSS and promoters but induces increased H3K4me3Q5dop more broadly, with peaks localizing to both genic and intergenic regions (Fig. **5e,f**). Motif finding analyses identified binding sites for homeobox transcription factors involved in brain development^42^, including GSX2, LHX1, and EMX2, enriched in both the upregulated H3K4me3Q5dop peaks and in the predicted promoters of the ZTR+sertraline-repressed genes from our RNA-seq results (Fig. **5g**). Finally, GO analysis on the genes found nearest to the differential H3K4me3Q5dop peaks further demonstrates that CDK9i+SSRI alters H3 dopaminylation at genes involved in kinase signaling, neuronal differentiation, nervous system development, and brain disorders (Fig. **5h,i**). Together, these results suggest that CDK9i+SSRI treatment regulates the transcription of genes related to neuronal signaling and development through a confluence of neurotransmitter-and CDK9i-mediated remodeling of H3 dopaminylation.

### Combining ZTR with SSRI induces distinct downstream kinase signaling events

CDK9 inhibition causes complex changes in protein phosphorylation^18,40,43–45^, and monoamine neurotransmitters similarly regulate various downstream kinases (e.g., PKA, PKC, CaMKII)^46^ via secondary messengers (cAMP, Ca^2+^) and through G protein-coupled receptor kinases (GRKs) and β-arrestin^47–49^. We next sought to characterize downstream kinase signaling mechanisms mediating DMG sensitivity to CDK9i+SSRI combination therapy. We find that 6 hours treatment of SU-DIPGVI cells with either ZTR alone or in combination with 5 µM sertraline reduces pS2-Pol2 (a CDK9 substrate), reduces pS10-H3 (a G2/M marker), and increases cl-CASP3 (an apoptotic marker) by western blotting (Fig. **6a**). Co-treatment with ZTR+sertraline results in a more pronounced decrease in pS2-Pol2 compared to ZTR alone (compare lanes 6-8 to 2-4; **Fig. 6a**). Treating BT245 cells with combinations of ZTR+duloxetine or ZTR+asenapine similarly leads to a stronger reduction in pS2-Pol2 than ZTR alone (compare lanes 2, 5, and 8; **Fig. 6b**), suggesting exacerbated changes in protein phosphorylation due to combination therapy.

**Fig. 6:**
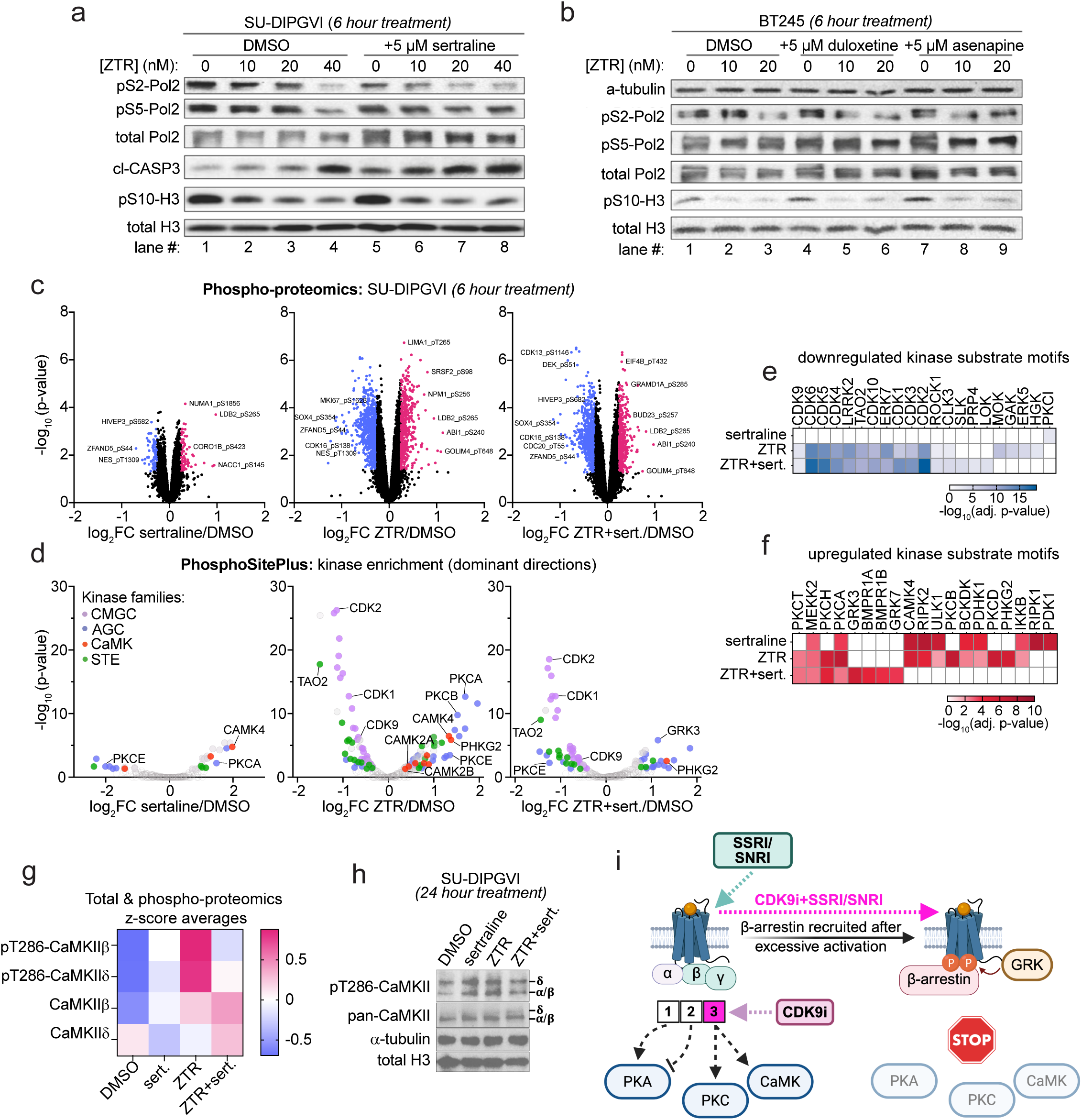
Combining ZTR with SSRI induces distinct downstream kinase signaling events. **a**,**b**, Western blot analysis of lysates from SU-DIPGVI cells (**a**) or BT245 cells (**b**) collected following 6 hours of treatment with ZTR (0, 10 nM, 20 nM) combined with vehicle (DMSO), sertraline (5 µM), duloxetine (5 µM), or asenapine (5 µM). **c**,**d**, Volcano plots of log_2_ fold change (log_2_FC) and-log_10_(adjusted *p*-values) from phospho-proteome profiling (**c**) and kinase motif enrichment analysis showing the dominant direction of differential phosphorylation (**d**) in SU-DIPGVI cells treated with vehicle (DMSO), 5 µM sertraline (sert.), 20 nM ZTR, or ZTR+sert for 6 hours. **e**,**f**, Consensus kinase motifs from the PhosphoSitePlus database that were either decreased (**e**) or increased (**f**) upon ZTR and/or sertraline treatment. **g**,**h**, Heatmap of z-scores from phospho-proteome and total proteome profiling (**g**) and Western blot analysis (24 hours treatment, **h**) on SU-DIPGVI cells exposed to DMSO, 5 µM sertraline (sert.), 20 nM ZTR, or ZTR+sert. showing altered CaMKII phosphorylation. **i**, Working model illustrating the effects of CDK9i+SSRI treatment on GPCR signaling to downstream kinases in DMG.

To more comprehensively assess the consequences of CDK9i+SSRI treatment on DMG signal transduction, we performed total-and phospho-proteomic analysis on SU-DIPGVI cells treated with vehicle, ZTR, sertraline, or a ZTR+sertraline drug combination (Fig. **6c,d** and Supplementary Table 6). Total-proteomic analyses show that ZTR+sertraline treatment upregulates proteins involved in RNA processing and Pol2 transcription compared to DMSO control (Extended Data Fig. **6a**) and reduces the overall protein abundance of monoamine signaling regulators and transcription factors previously linked to glioma progression (SOX9, ZFP36L1^50,51^; Extended Data Fig. **6a,b**). Our phospho-proteomic profiling identified many differential phosphorylation sites following ZTR or ZTR+sertraline treatment relative to vehicle control (adjusted *P*<0.05, log2FC+/-0.4; Supplementary Table 6), including known CDK9-binding partners (curated from STRING and BioGRID; Extended Data Fig. **6c**), verifying a disruption in CDK9-associated protein phosphorylation events.

GO analyses of these phospho-proteomic data reveal a partial overlap between the downstream targets of ZTR, sertraline, and ZTR+sertraline combination treatment, including proteins linked to RNA regulation and Rho GTPase signaling (Extended Data Fig. **6d-f**). Kinase motif enrichment analysis suggests that either ZTR monotherapy or ZTR+sertraline treatment reduces the phosphorylation of CMGC family kinase substrates (CDK, MAPK, GSK, CDLK), including known CDK9 targets (Fig. **6d,e**), and reveals increased phosphorylation of CaMK and AGC family kinase substrates targeted by kinases like CaMKII and PKC (Fig. **6d,f**). In contrast, ZTR+sertraline co-treatment did not induce a significant change in CaMKII-or PKC-targeted motif activity but rather increased GPCR kinase (GRK) substrate phosphorylation (Fig. **6d,f**).

We selected altered CaMKII phosphorylation for further validation based on known roles for CaMK signaling in gliomagenesis and glioma stem cell maintenance^52,53^. In our phospho-proteomic analysis and by western blotting, both sertraline and ZTR treatment increased autophosphorylation of the T286 residue of CaMKII (pT286-CaMKII), a marker of constitutive activation^46^, while ZTR+sertraline had minimal effect on this phosphosite (Fig. **6g,h**). These results suggest that SSRI or CDK9i treatment alone activates CaMK family kinases in DMG, but this aberrant pattern of kinase activation is prevented by SSRI and CDK9i combination treatment. Based on these data, we constructed a working model of CDK9i+SSRI/SNRI synergy where treating DMG cells with sertraline or ZTR alone induces G-protein-biased GPCR kinase signaling cascades to activate CaMKII, whereas ZTR+sertraline treatment causes excessive levels of GPCR signaling, leading to negative feedback inhibition of neurotransmitter receptors by GRK/β-arrestin^47^ (Fig. **6i**).

### Neurotransmitter GPCR agonists and antagonists synergize with CDK9i by reducing CaMKII activity

Our working model of CDK9i+SSRI-induced negative feedback inhibition of GPCRs by GRK/β-arrestin (Fig. **6g**) affords a possible explanation for the paradoxical identification of neurotransmitter GPCR antagonists and SSRI/SNRI, which increase monoamine neurotransmitter signaling, as potent CDK9i-synergizers (Fig **3a,b** and Supplementary Table 3). To assess our model of CDK9i synergy with neurotransmitter modulation in DMG, we next determined potential drug synergy between ZTR and additional classes of neuropsychiatric drugs. Co-treatment with ZTR and neurotransmitter GPCR agonists, including the D2 dopamine receptor (D2R) agonists rotigotine (RTG) and cabergoline, led to a synergistic reduction in the growth of DMG cells (SJ-DIPGX37; Fig. **7a** and Extended Data Fig. **7a**), mirroring our findings with ZTR+SSRI/SNRI (Fig. **3** and Extended Data Fig. **3a-d**). ZTR similarly synergizes with UNC9994, a tool compound that activates β-arrestin-biased D2R signaling (Fig. **7a** and Extended Data Fig. **7a**). DMG cells were also sensitive to combinations of ZTR and antagonists against dopamine, serotonin, and norepinephrine receptors, including mirtazapine, olanzapine, silodosin, and nebivolol, which inhibit GPCR signaling (Fig. **7b** and Extended Data Fig. **7b**). Finally, we find that trifluoperazine (TFP), a dual antagonist of both D2R and CaM, synergistically reduced DMG growth in combination with ZTR (Fig. **7b** and Extended Data Fig. **7b**), consistent with our finding that ZTR+sertraline treatment prevents CaMKII activation by ZTR treatment (Fig. **6e-h**). We analyzed pT286-CaMKII in SJ-DIPGX37 cells treated for 24 hours with vehicle, RTG (10 µM), TFP (2 µM), and ZTR (20 nM), and find that both the GPCR agonist combination (ZTR+RTG) and the GPCR antagonist combination (ZTR+TFP) reduced pT286-CaMKII relative to ZTR alone (compare lanes 3 to 4 and 7 to 8; Fig. **7c**), as predicted. These results further support our model whereby neurotransmitter-targeting drugs prevent CDK9i-induced CaMKII activation through convergent signaling mechanisms.

**Fig. 7:**
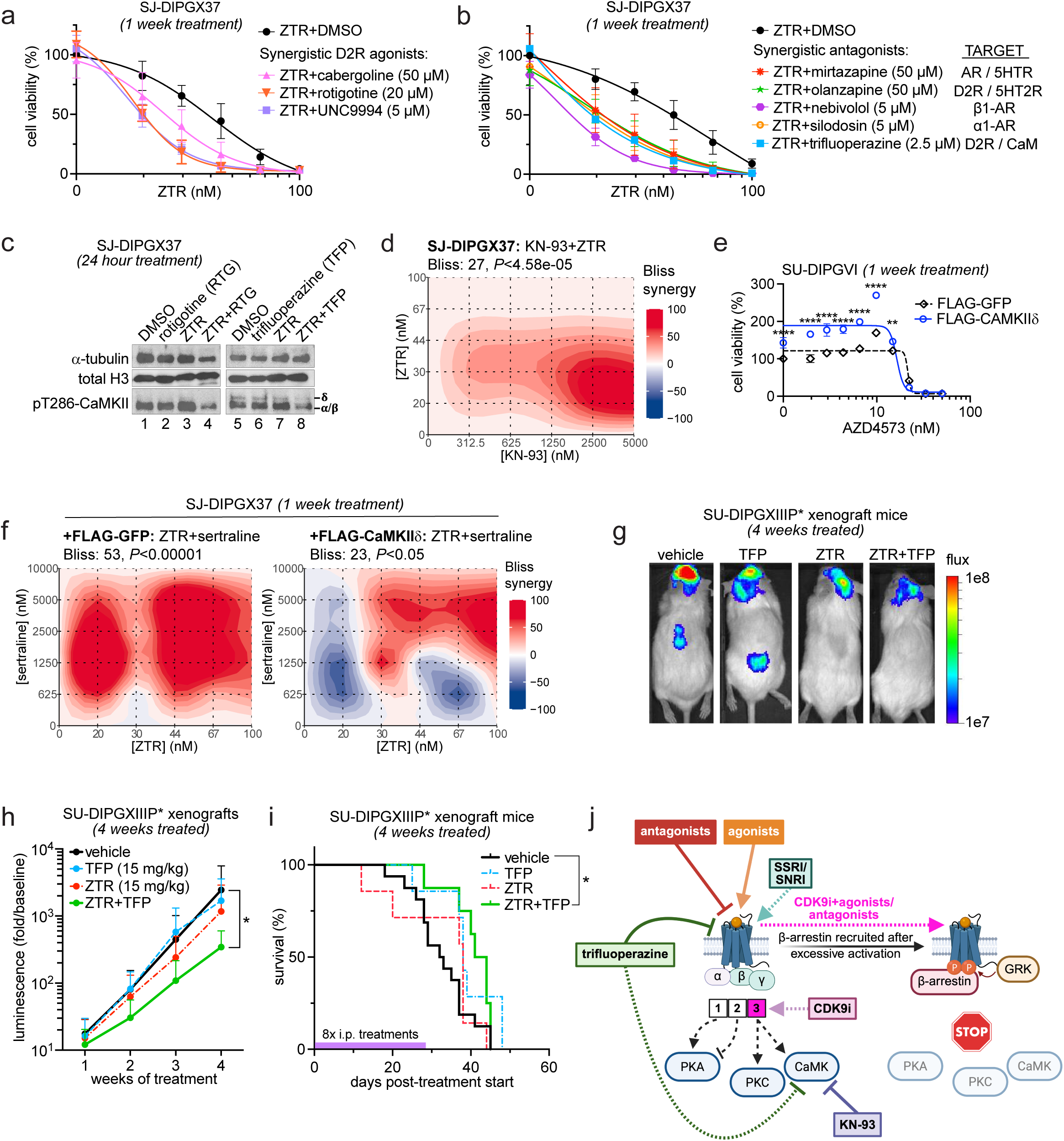
Neurotransmitter GPCR agonists and antagonists synergize with CDK9i by reducing CaMKII activity. **a**,**b**, Dose curves showing SJ-DIPGX37 cell viability (CellTiter-Glo) after 7 days of treatment with ZTR in combination with vehicle (DMSO) or neurotransmitter GPCR antagonists (**a**) and agonists (**b**). For **a**,**b**, *n* = 3 replicates per group. **c**, Western blot analysis of SJ-DIPGX37 whole cell lysates collected following 24 hours of treatment with DMSO, RTG (10 µM), TFP (2 µM), ZTR (20 nM), ZTR+RTG, or ZTR+TFP. **d**, Bliss synergy score heatmap of cell viability (CellTiter-Glo) for SJ-DIPGX37 after 7 days of treatment with dose matrices of ZTR and the CaM inhibitor, KN-93. **e**, Dose curves showing the effect of 7 days of treatment with AZD4573 on the viability (CellTiterGlo) of SU-DIPGVI cells overexpressing either FLAG-tagged CaMKIIδ or GFP control. For **3**, *n* = 3 replicates per group. **f**, Bliss score heatmaps showing ZTR+sertraline drug synergy in SJ-DIPGX37 cells transduced with FLAG/HA-tagged GFP (left panel, negative control), FLAG-tagged CaMKIIδ (right panel). For the FLAG/HA-GFP group, mean Bliss score = 53 & *P*<0.00001, and for the FLAG-CaMKIIδ group, mean Bliss score = 23 & *P*<0.05. **g**,**h**, Representative images (**g**) and quantification of fold change in BLI signal (**h**) in SU-DIPGXIIIP* xenograft mice treated for 2 weeks with vehicle, ZTR (15 mg/kg), TFP (15 mg/kg), or ZTR+TFP. **i**, Kaplan-Meier survival analysis indicating that treatment with a combination of ZTR and TFP significantly increased survival in the SU-DIPGXIIIP* xenograft model (purple bar shows treatment period; *P*<0.0327, Gehan-Breslow-Wilcoxon test). For **h**,**i**, *n* = 16 mice for vehicle group, *n* = 7 mice for the ZTR group, *n* = 7 mice for the TFP group, and *n* = 8 mice for ZTR+TFP group. **j**, Working model showing how SSRI/SNRI and neurotransmitter GPCR antagonists and agonists synergize with CDK9 inhibitors by reducing aberrant activation of CaMK family kinases.

We conducted additional studies to determine whether CaMK modulation is sufficient to regulate DMG response to CDK9i. Indeed, co-treatment with a CaM inhibitor (KN-93) and ZTR resulted in a synergistic reduction in SJ-DIPGX37 cell growth (Fig. **7d**). In contrast, overexpressing a FLAG-tagged isoform of CaMKII (FLAG-CaMKIIδ) induced a 1.5-fold increase in SU-DIPGVI cell growth even in the presence of low concentrations of the CDK9i AZD4573 (10 nM) compared to SU-DIPGVI cells overexpressing FLAG/HA-GFP as a control (Fig. **7e** and Extended Data Fig. **7c**), suggesting that CaMKII signaling promotes DMG growth. Overexpression of CaMKIIδ also dampened ZTR+sertraline drug synergy in SJ-DIPGX37 compared to GFP control (Fig. **7f** and Extended Data Fig. **7d**). These findings identify CaMKII signaling as a key determinant of drug synergy between neurotransmitter-targeting agents and CDK9i in DMG.

We selected TFP for follow-up *in vivo* testing since it synergizes strongly with ZTR (Extended Data Fig. **7b**) and may be suitable for repurposing as a treatment for glioblastoma since it is FDA-approved for schizophrenia and anxiety^54^. We tracked tumor growth and overall survival in SU-DIPGXIIIP* and SJ-DIPGX37+ZsGreen/luciferase xenograft mice treated with vehicle, TFP, ZTR, or ZTR+TFP (15 mg/kg each drug, 2 days/week for 4 weeks; Fig. **7g-i** and Extended Data Fig. **7e**). While both ZTR and ZTR+TFP reduced the SU-DIPGXIIIP* xenograft BLI signal (Extended Data Fig. **7e**), only ZTR+TFP co-treatment significantly reduced SJ-DIPGX37 xenograft growth (Fig. **7g,h**). ZTR+TFP co-treatment also improved survival of the SJ-DIPGX37 xenograft mice compared to vehicle control (Fig. **7i**). These results support a model where SSRI/SNRIs and neurotransmitter GPCR agonists and antagonists synergize with ZTR to reduce DMG growth through GPCR-CaMK pathway inhibition (Fig. **7j** and Extended Data Fig. **7f**).

## Discussion

RNA polymerase II (Pol2) transcriptional dysregulation and aberrant neurotransmitter signaling are key drivers of gliomagenesis and represent actionable vulnerabilities that can be modulated using Pol2-and neurotransmitter-targeting agents^3,10^. The discovery of histone monoaminylation^15^ raises the possibility that neurotransmitters may directly influence chromatin and transcriptional regulation in brain tumors, including H3K27M mutant diffuse midline glioma (DMG), a disease characterized by profound epigenetic dysfunction. However, how monoamine neurotransmitter signaling influences aberrant chromatin and gene regulatory states driving DMG tumor growth and response to therapy remains poorly understood^10,55^.

H3K27M oncohistones induce widespread chromatin and transcriptional reprogramming, including the dysregulation of neurodevelopmental and glial lineage genes^26,28^. Histone monoaminylation at H3Q5 via dopamine and serotonin (H3Q5dop and H3Q5ser) has been implicated in the regulation of neuronal differentiation, stress responses, reward circuitry, and malignancy in other biological contexts^17,56^, but potential functions of histone monoaminylation have not yet been reported in DMG to the best of our knowledge. Our chromatin profiling of histone dopaminylation in DMG cells reveals that H3K4me3Q5dop-dominant genes exhibit high pS2-Pol2 and H3K27M occupancy and encode Pol2 transcriptional cofactors, whereas H3K4me3Q5dop-low genes preferentially encode neuronal and synaptic signaling regulators. These findings suggest that functionally distinct gene sets exhibit divergent H3 dopaminylation patterns in DMG. This complex interplay between H3K27M and H3K4me3Q5dop implicates histone monoaminylation in the aberrant transcription of neurodevelopmental genes and Pol2 cofactors central to DMG pathogenesis.

Although altered neurotransmitter signaling has been linked to glioma development and progression, little is known about the impact of cell-intrinsic or local neurotransmitter signaling on DMG response to therapy. We observe differential expression of neuronal signaling genes in DMG xenograft tumors exhibiting resistance to the Pol2-targeting CDK9 inhibitor (CDK9i) zotiraciclib (ZTR). Consistent with this observation, pharmacologic modulation of monoamine neurotransmitter signaling strongly synergized with CDK9 inhibition to reduce DMG cell viability and suppress tumor growth in orthotopic xenografts. Together, these results demonstrate that neurotransmitter signaling regulates DMG response to CDK9i and that co-treatment with FDA-approved drugs to modulate monoamine neurotransmitter signaling may overcome resistance to CDK9-targeted therapies.

Further integrated chromatin and transcriptomic profiling studies reveal that synergistic CDK9i+SSRI combination treatment remodels chromatin accessibility and alters both pS2-Pol2 occupancy and H3 dopaminylation at neurodevelopmental and synaptic genes, accompanied by robust changes in gene transcription. These dynamic chromatin changes were evident at the TSS of genes like *GRIA4*, which encodes a glutamate receptor that was upregulated by chronic CDK9i treatment *in vivo* and suppressed by CDK9i+SSRI combination therapy. Consistent with these observations, a recent paper similarly found that another CDK9i (atuveciclib) increased Pol2 pausing and altered the transcription of neuronal signaling genes in DMG^3^. CDK9i+SSRI treatment also altered H3 dopaminylation within DMG bivalent H3K4me3+/H3K27me3+ promoters, suggesting that histone dopaminylation may represent a previously unrecognized mechanism modulating developmental chromatin transitions and transcriptional plasticity in DMG.

Additional studies aimed at uncovering the impact of CDK9i+SSRI on DMG kinase signaling reveal that CDK9 inhibition drives pro-survival CAMKII activation, whereas the addition of SSRIs or neurotransmitter GPCR agonists or antagonists in combination with CDK9i suppressed CAMKII signaling. We propose a working model in which both neurotransmitter agonists and antagonists paradoxically synergize with CDK9i through convergent mechanisms, leading to CAMKII inactivation and growth suppression. Potential mechanisms connecting treatment-induced changes in GPCR activity and Ca^2+^/CaMK signaling to altered H3 dopaminylation patterns and chromatin state in DMG will be an important area for future investigation.

Finally, in orthotopic xenograft models, co-treatment with CDK9i and either an antidepressant (sertraline) or an antipsychotic (trifluoperazine) significantly improved overall survival. Several of the CDK9i-synergistic drugs identified in our cell culture-based assays, such as olanzapine and escitalopram, are commonly prescribed to pediatric brain tumor patients to alleviate symptoms like nausea, anxiety, and depression, suggesting that these drugs might be repositioned to enhance DMG response to CDK9 inhibitors or other drugs targeting transcriptional regulation. Compounds targeting monoamine neurotransmitter signaling inhibit solid tumor growth^57–60^, and SSRIs were recently found to improve immunotherapeutic responses in other cancers^60^. Co-targeting CDK9, neuronal signaling pathways, and other known vulnerabilities may therefore provide new therapeutic opportunities to improve outcomes for patients with DMG and other brain tumors.

## Data Availability Statement

Data will be made available upon reasonable request. Sequencing data generated in this study is available on the Gene Expression Omnibus (GEO) under accession number GSE324146. Published H3K27M and H3K27me3 profiling data are available on GEO under accession number GSE162976.

## Methods

### Ethics statement

The Institutional Animal Care and Use Committee at Baylor College of Medicine approved all animal protocols.

### Cell culture

The patient-derived H3K27M DMG cell lines (SU-DIPGXIII, SU-DIPGIV, SU-DIPGVI, BT245, SJ-DIPGX37, and SU-DIPGXIIIP*) used in this study were kindly provided by Dr. Michele Monje, Dr. Suzanne Baker, and Dr. Keith Ligon, while the adherent control cell lines (NHA-hTERT and BJ-Fibroblasts) and HEK293T were purchased from ATCC. All DMG cells were grown in suspension as neurospheres in serum-free glioma stem cell culture media containing a 50:50 mix of DMEM/F-12 (cat# 10-092-CV, Corning) and Neurobasal-A (cat# 10888022, Gibco) supplemented with B27 minus vitamin A (cat# 12587010, Gibco), 1 mM sodium pyruvate (cat# 25000CI, Corning), 2 mM GlutaMAX (cat# 35050079, Gibco), 1x MEM non-essential amino acids (cat# 11140050, Gibco), 0.01 M HEPES (cat# 25-060-CI, Corning), Antibiotic-Antimycotic (cat# 30-004-CI, Corning), 0.0002% Heparin (cat# 07980, StemCell Technologies), hPDGF-AA (20 ng/mL; cat# 10016100UG, Fujifilm Irvine Scientific Inc), hPDGF-BB (20 ng/mL; cat# 1001810, Fujifilm Irvine Scientific Inc), hEGF (20 ng/mL; cat# 100261MG, Fujifilm Irvine Scientific Inc), and hFGF154 (20 ng/mL; cat# 100146500UG, Fujifilm Irvine Scientific Inc). Cells were maintained by passaging every 4-6 days using TrypLE Express (cat# 12604021, ThermoFisher). The adherent cell lines were cultured in DMEM (cat# MT10013CV, Corning) containing 10% FBS (cat# A5670701, Fisher Scientific) and 1% penicillin/streptomycin (cat# SV3007901, Fisher Scientific) and were passaged using 0.25% trypsin/EDTA (cat# MT25053Cl, Fisher Scientific). All cell lines were grown in a 5% CO_2_ incubator at 37°C.

### Cell transfections and infections

Lentivirus packaging was done by co-transfecting HEK293T cells with a plasmid of interest (pHIV-Luc-Zs-green or plenti-FLAG-CAMK2D), pCMV-dR8.2-dvpr (cat# 8455, Addgene), and pCMV-VSV-G (cat# 8454, Addgene) in a 10-cm dish (1 μg each) using PEI (cat# 764965, MilliporeSigma). Secreted virus was syringe-filtered (0.45-μm) from the medium 36-48 hours post-transfection, then mixed with 50% PEG8000 (cat# BP233-1, Fisher Scientific) and incubated overnight at 4°C. Viral supernatants were then concentrated by centrifugation at 960xg for 20 minutes and resuspended in 600 µl of neurosphere culture media. For lentiviral infections, 150 µl of concentrated lentivirus was combined with 1e6 DMG cells in 6 wells of a 12-well plate, then centrifuged at 960xg for 90 minutes at room temperature and transferred to cell culture flasks to recover overnight in a 5% CO_2_/37°C incubator. After 48-72 hours, cells were either sorted by FACS based on positive fluorescent protein expression or selected with puromycin (2 µg/mL; cat# sc-108071A, SantaCruz Biotechnology) as appropriate. For xenograft studies, SU-DIPGXIIIP* cells were infected with a ZsGreen/luciferase lentiviral construct (pHIV-Luc-ZsGreen was a gift from Bryan Welm; cat# 39196, Addgene), and for cell viability assays, SU-DIPGVI cells were infected with flag-tagged GFP or CaMKIIδ (plenti-FLAG-CAMK2D was a gift from Derrick Sek Tong Ong; cat# 221699, Addgene).

### Targeted neurotransmitter metabolomics

DMG cells were dissociated and washed with PBS, then biological triplicates of 5e6 cells for each cell line were pelleted and stored at-80°C prior to processing. Thawed cell pellets were then lysed by sonication (2 pulses, 30 seconds each) in a 50:50 mixture of water and methanol. Proteins were removed by chloroform extraction and filtration (Amicon Ultracel 3-kDa Membrane; cat# UFC9003, MilliporeSigma) before the samples were concentrated by speed vacuum. Targeted metabolomics for neurotransmitters was performed on an Agilent (6490 QQQ triple quadrupole) and SCIEX TOF mass spectrometers. Metabolites were targeted with electrospray source ionization voltages (+4,000 V and-3,500 V) using multiple reaction monitoring and quantified against internal standards.

### Cell viability and CDK9i synergy assays

Cell viability was either assessed by reading CellTiter-Glo luminescence on a Perkin-Elmer plate reader or by manual live cell counts using a hemacytometer and the Trypan Blue (cat# MT25900CI, Fisher Scientific) exclusion method. For the CDK9i drug synergy screen, drugs from the Tocriscreen FDA-approved Drugs Library were plated into white 384-well plates with an Echo Acoustic Liquid Handler, followed by plating 2,500 cells in neurosphere culture media containing either vehicle (DMSO) or 40 nM ZTR and then performing CellTiter-Glo assays after 7 days treatment. For validation studies, 2,500-10,000 DMG cells/well were plated in 384-or 96-well plates containing drug combinations, and viability was similarly assessed using CellTiter-Glo after 7 days. We determined drug synergy by Bliss analysis and generated heatmaps of synergy scores using the synergyfinder R package (version 3.12.0) with the default settings.

### Immunofluorescence staining

Histone dopaminylation and NET expression were visualized in DMG cells plated on laminin-coated coverslips in a 12-well plate with 1.5e5 cells per well, fixed with 4% paraformaldehyde in PBS for 20 minutes, washed with PBST, blocked for 24 hours (5% FBS, 0.3% triton, 1XPBS), and stained overnight at 4°C with anti-H3K4me3Q5dopaminyl (cat# ABE259010, MilliporeSigma) or anti-NET (cat# NBP3-12251, Novus Biologicals) diluted to 1:500 in antibody solution (1% BSA, 0.3% Triton-X 100, 1XPBS) with gentle rocking. Coverslips were then washed with PBST 3 times and incubated with an Alexa-fluorophore conjugated secondary antibody diluted to 1:500 in antibody solution for 30 minutes at room temperature, then washed with PBST 3 times and stained with Hoechst at a 1:1,000 dilution for 15 minutes. Coverslips were mounted on microscope slides with Prolong Gold Antifade reagent, and cells were imaged using a Keyence BZ-X800 microscope.

### Western blotting

Whole cell lysates were prepared by resuspending cell pellets in RIPA lysis buffer (50 mM Tris-HCl pH 8.0, 0.1% SDS, 1% NP-40, 0.5% sodium deoxycholate, and 150 mM NaCl) with protease inhibitors (cat# 5892970001, MilliporeSigma), and phosphatase inhibitors (cat# 4906837001, Millipore Sigma). Samples were then prepared in 1xLaemmli buffer with 2.5% v/v β-mercaptoethanol (cat# 97064-878, VWR) and stored at-80°C. Lysates were run on 4-20% SDS-PAGE gels (cat# 4561096, BioRad) and transferred to nitrocellulose membranes (cat# 10600001, Cytiva) on ice at 100 V for 35 minutes. Membranes were then blocked in 5% BSA in PBST for 15 minutes at room temperature before overnight incubation at 4°C with the following primary antibodies diluted in 5% BSA: total histone H3 (1:500,000; cat# ab1791, Abcam), α-tubulin (1:5,000; cat# A11126, ThermoFisher), SERT (1:1,000; cat# ab102048, Abcam), DAT (1:1,000; cat# MA5-24796, ThermoFisher), pS2-Pol2 (1:1,000; cat# 04-1571, MilliporeSigma), pS5-Pol2 (1:1,000; cat# 04-1572, MilliporeSigma), total Pol2 (1:1,000; cat# 502075139, Fisher Scientific), cl-CASP3 (1:1,000; cat# 9661S, Cell Signaling Technology), pS10-H3 (1:5,000; cat# 9701S, Cell Signaling Technology), pT286-CaMKII (1:1,000; cat# 12716S, Cell Signaling Technology), and pan-CaMKII (1:1,000; cat# 4436T, Cell Signaling Technology). The next day, membranes were washed 3 times for 5 minutes with PBST then incubated with 1:10,000 anti-rabbit-HRP (cat# B40962, Invitrogen) or anti-mouse-HRP (cat# B40961, Invitrogen) secondary antibodies in 5% milk in BSA for 45 minutes at room temperature. Membranes were washed 3 times for 5 minutes with PBST then imaged on a Biorad ChemiDoc using ECL (cat# A38555, ThermoFisher).

### Xenograft mouse models and treatments

DMG tumors were generated by stereotactic injection of 2.5e5 or 5e5 SU-DIPGXIIIP*+ ZsGreen/luciferase cells into the pons of 6-to 12-week-old NSG mice (NOD.Cg-Prkdc^scid^Il2rg^tm1Wjl^/SzJ) as previously described^61^. In Figure **2a-c**, mice were treated 4 consecutive days per week for 4 weeks by oral gavage (p.o.) with vehicle (7% DMSO, 0.5% (w/v) methylcellulose, and 0.1% Tween 80 in saline) or 15 mg/kg ZTR (zotiraciclib). In Figure **3e-g** and Extended Data Figure **3f,g**, treatment consisted of vehicle (7% DMSO in saline), 2.5 mg/kg AZD4573, 2.5 mg/kg sertraline, or AZD4573+sertraline dosed by intraperitoneal (i.p.) injection every other day for 3 days per week for 4 weeks. In Figure **3h-k**, mice were treated by i.p. injection with vehicle (7% DMSO, 40% PEG300, and 1% Tween 80 in saline), 20 mg/kg ZTR, 5 mg/kg duloxetine, or ZTR+duloxetine every other day for 3 days per week for 2 weeks. Finally, in experiments shown in Figure **7g-i** and Extended Data Figure **7f,** mice were treated by i.p. injection with vehicle (4% DMSO, 40% PEG300, and 1% Tween 80 in saline), 15 mg/kg ZTR, 15 mg/kg trifluoperazine (TFP), or ZTR+TFP once every 3-4 days over 4 weeks for a total of 8 treatments. Mice treated with TFP or ZTR+TFP were monitored for side effects, including slow movement and gastrointestinal issues, which resolved within 24 hours after treatment. Bioluminescent imaging was done on isoflurane-anesthetized mice ∼10 minutes after i.p. injection of D-luciferin (150 mg/kg) diluted in saline using an IVIS X5.

### ZTR-resistant xenograft tumor isolation

Brains were dissected from DIPGXIIIP* xenograft mice treated with vehicle or ZTR (15 mg/kg, 3 times per week) for 3 weeks. Treated mice were euthanized with 3 L/min CO_2_ for 3 minutes and perfused with ice-cold 1X PBS prior to brain collection. A mouse brain dissociation kit (cat# 130-110-201, Miltenyi Biotec) and debris removal solution (cat# 130-109-398, Miltenyi Biotec) were used to obtain a single cell suspension, then mouse cells were depleted using a mouse cell depletion kit (cat# 130-104-694, Miltenyi Biotec) and an AutoMACS.

### Immunohistochemistry (IHC)

Pol2 phosphorylation at serine 2 (pS2-Pol2) was quantified in the pons of SU-DIPGXIIIP* xenograft mice treated with vehicle, AZD4573, sertraline, or AZD4573+sertraline for 3 weeks. Brains were collected approximately 24 hours after the last treatment from mice euthanized and perfused with ice-cold 1X PBS followed by 4% paraformaldehyde. Isolated brains were post-fixed in 4% PFA for 3 hours and then transferred to sucrose solution (30% sucrose, 1X PBS) for 24 hours then embedded in OCT and stored at-80°C. Frozen sections (12 µm, coronal) were placed onto microscope slides and stained with anti-pS2-Pol2 (1:1,000ic; cat# 04-1571, MilliporeSigma), washed with PBST, stained with a secondary Alexa-fluorophore conjugated antibody, then incubated with Hoechst (1:1,000) and mounted with a coverslip using Prolong Gold Anti-fade reagent. For BrdU staining experiments, mice received a 2 hour pulse of BrdU (100 mg/kg) administered 24 hours after the final drug treatment followed by PFA perfusion as described above, post-fixation in 4% PFA overnight, and storage in 70% ethanol. Brain tissues were paraffin-embedded, sectioned and processed for IHC using anti-BrdU (clone ZBU30; cat# 03-3900, Invitrogen) and anti-cl-CASP3 (cat# 9661, Cell Signaling Technologies) antibodies with the assistance of the Smith Breast Center Pathology lab at Baylor College of Medicine using standard protocols on the Shandon Varistain (24-4) Automatic Stainer. Slides were imaged using a Keyence BZ-X800 microscope and representative images were analyzed using QuPath.

### Analysis of pediatric glioma patient proteomics datasets

Pediatric brain tumor proteomic data were sourced from the PedcBio data portal (CPTAC provisional proteomics dataset, https://pedcbioportal.org/) and a ranked Spearman’s correlation analysis for CDK9 protein expression in each sample was performed against the expression of all other detected proteins.

### RNA-seq

RNA-seq was done on biological triplicates of SU-DIPGXIIIP* tumor cells isolated from xenografted NSG mice treated with vehicle or ZTR (15 mg/kg, 3 times per week, p.o.) for 3 weeks or performed in triplicate samples of 2.5e5 cultured SU-DIPGXIII cells dissociated after 24 hours treatment with DMSO, ZTR (20 nM), AZD4573 (6 nM), sertraline (5 µM), ZTR+sertraline, or AZD4573+sertraline. RNA was extracted with TRIzol reagent (cat# 15596026, ThermoFisher) and column-purified (cat# 74104, Qiagen) before cDNA generation from polyA RNA. Sequencing libraries were prepared from cDNA using Illumina protocols, followed by paired-end sequencing on an Illumina NovaSeq 6000. Reads were aligned to the genome with STAR (hg38), and differential gene expression was analyzed with DESeq2.

### CUT&RUN

CUT&RUN was done in duplicate for pS2-Pol2, H3K4me3, and H3K4me3Qdop on 5e5 untreated SU-DIPGXIII cells and on 5e5 SU-DIPGXIII cells treated for 24 hours with DMSO or a combination of ZTR (20 nM) and sertraline (5 µM). Neurospheres were dissociated to single cells with TrypLE Express Enzyme, washed with 1X HBSS, counted, and prepared for CUT&RUN with 500,000 cells/sample. CUT&RUN-enriched DNA was prepared using the Cell Signaling Technologies CUT&RUN Kit (cat# 86652) following the manufacturer’s protocol using the following antibodies: pS2-Pol2 (cat# 04-1571, MilliporeSigma), H3K4me3 (cat# C15410003, Diagenode), H3K4me3Q5dop (Anti-H3K4me3Q5dopaminyl; cat# ABE2590, MilliporeSigma), and IgG (Anti-IgG Rabbit; cat# C15410206, DIAgenode). Yeast spike-in DNA (S. cerevisiae; Cell Signaling Technology) was added at 0.5% to each reaction following chromatin digestion. Sequencing libraries were prepared from 3 ng of enriched DNA using the SMARTer ThruPLEX Plasma-seq kit (cat# R400681, Takara Bio) with dual indexes (cat# DX34752, Takara Bio) according to the manufacturer’s protocol. Adapter-ligated libraries were amplified for 11 PCR cycles, purified using AMPure XP beads (Beckman Coulter), and evaluated for fragment size distribution on an Agilent 2100 Bioanalyzer. Library concentrations were determined by quantitative PCR with the KAPA Library Quantification Kit (Roche) before pooling for sequencing on an Illumina NovaSeq 6000 to generate 150 bp paired-end reads. Reads were processed with trimgalore and aligned to the genome (hg38) with bowtie2, then peaks were called with MACS2 followed by differential peak analysis with diffbind.

### ATAC-seq

ATAC-seq was performed in triplicate on 5e5 SU-DIPGXIII cells treated for 24 hours with DMSO or a combination of ZTR (20 nM) and sertraline (5 µM) as previously described^62^. Briefly, DNA was tagmented with Tn5, and then purified for sequencing library preparation using Illumina protocols, followed by paired-end sequencing on an Illumina NovaSeq 6000. Reads were processed with TrimGalore and aligned to the genome with BWA (hg38). Peaks were then called with MACS2, and differential peak analysis was done with csaw.

### Transcription factor motif analysis

HOMER, with repeat-masking and the size parameter set to 200 bp around the center of peaks, was used to find enriched motifs in the differential ATAC-seq, pS2-Pol2, and H3K4me3Q5dop datasets and in the predicted promoters of genes of interest (8 to 10 bp motifs,-400 to +100 relative to the TSS).

### Total and phospho-proteomics

SU-DIPGVI cells were treated for 6 hours with DMSO, ZTR (20 nM), sertraline (5 µM), or ZTR+sertraline followed by total and phospho-proteomic analysis of triplicate samples of 1.5e7 cells. Quantitative mass spectrometry was performed using a TMT protocol and Thermo Fisher Orbitrap mass spectrometers. PhosphoSite Plus was used to analyze kinase substrate motif enrichment.

### Statistical analysis and graphics

GraphPad Prism 10.3 software was used to perform 2-way ANOVA and log-rank tests. Significance is reported after correction for multiple comparisons and plots show means ± standard deviations unless otherwise specified in the figure legend. Model figures were created in BioRender by R.L.M.

### Author Contributions

R.L.M. and J.N.A. designed and conceived the study, conducted bioinformatic analyses, performed experiments, and wrote the manuscript. B.R.E. performed xenograft, Western blot, and immunofluorescence analyses. R.U.R. and E.I.C. assisted with *in vitro* and *in vivo* studies. A.N.N. and N.T. conducted dose curve experiments. A.E. and C.M.O. provided drug libraries and helped set up drug screens. J.R.T. and A.K.D. assisted with mouse treatments and writing the manuscript. R.M. assisted with Western blotting. K.P. and D.C.K. optimized, prepped, and sequenced CUT&RUN libraries. K.Y. and B.D. assisted with study design and xenograft models.

A.S.H. pre-processed sequencing data sets and assisted with bioinformatic analyses.

## Acknowledgments

These studies were supported by the National Institutes of Health (NIH) (5R01NS129860 to J.N.A.), the ChadTough Defeat DIPG Foundation, the Matthew Larson Foundation (Iron Matt), and a research gift from the Houston Men of Distinction. Dr. Rebecca Murdaugh received training support from the Baylor College of Medicine (BCM) Center for Cell and Gene Therapy (CAGT) training grant from the NIH (5T32HL092332 to R.L.M.). Jack Tremblay and Alfred Dei-Ampeh received additional support from the BCM Training Program in Cell and Molecular Biology through the NIH (T32GM136560 to J.R.T. and A.D.). We thank the Genomics and RNA Profiling Core at Baylor College of Medicine for their assistance with CUT&RUN and RNA-seq studies, supported by the NIH (NCI, P30CA125123) and the Cancer Prevention and Research Institute of Texas (CPRIT, RP250580). ATAC-seq data were generated by the UTHealth Houston Cancer Genomics Core, supported by CPRIT (RP240610) and an NIH S10 grant (1S10OD036427). Additional support for this project was provided by the BCM CPRIT Proteomics and Metabolomics Core Facility, funded by CPRIT (RP210227), the NIH (P30CA125123), and the Dan L. Duncan Comprehensive Cancer Center. We also thank the Mouse Metabolism and Phenotyping Core, supported by the NIH (UM1HG006348, R01DK114356, R01HL130249), and the Smith Breast Center Pathology Lab for their assistance with live animal imaging and histopathological studies. This project was also supported by the BCM Cytometry and Cell Sorting Core, supported by CPRIT (RP240432) and the NIH (CA125123 and ODO36336).

