## Supplementary material for "H3 dopaminylation and CaMKII modulate diffuse midline glioma response to CDK9 inhibition": Figures S1-S7, Supplementary Table Legends

### Extended Data Figure 1

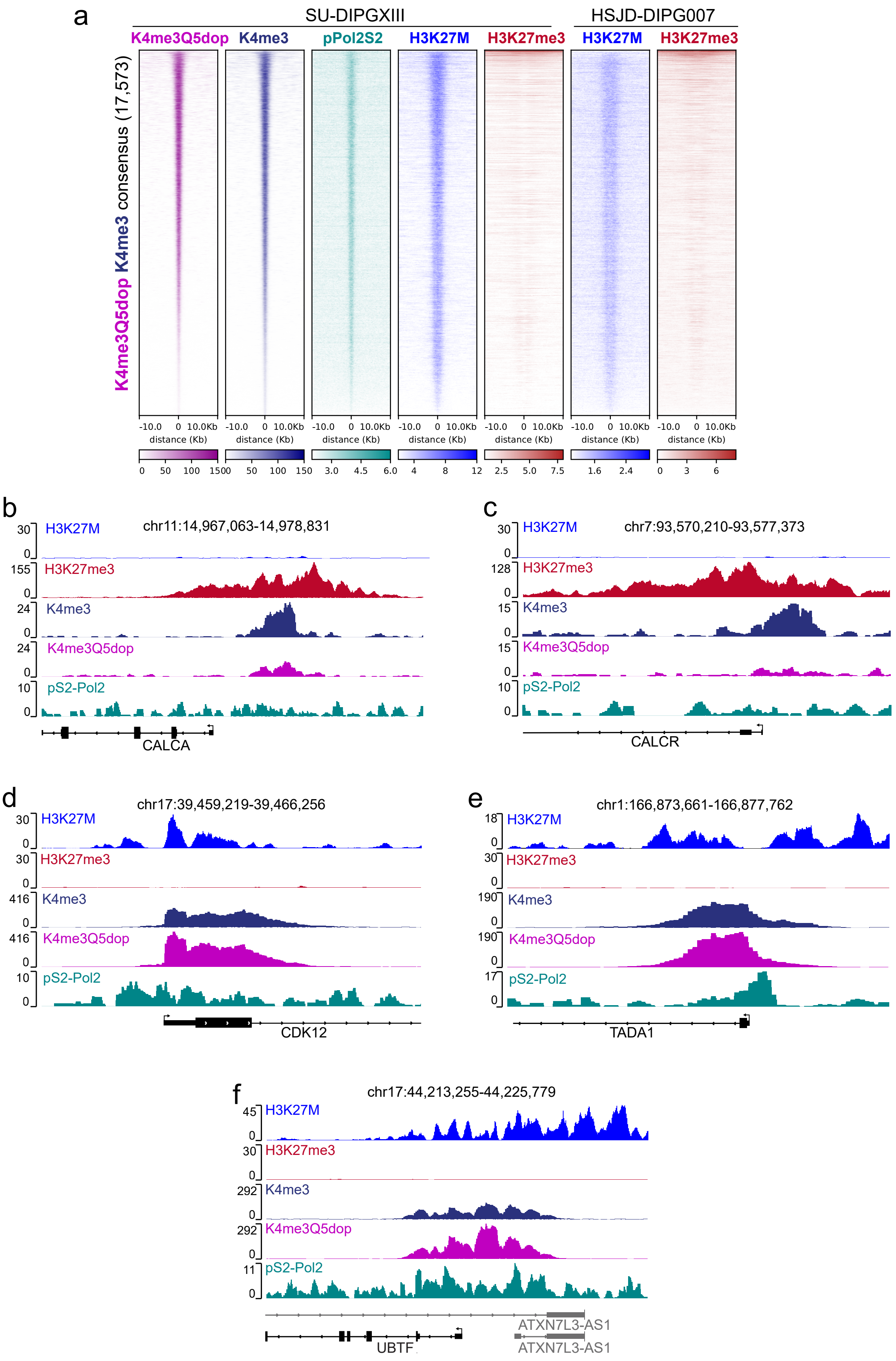

**Extended Data Fig. 1: H3 dopaminylation marks genes encoding Pol2 transcriptional regulators in DMG. a**, Heatmap showing consensus set of H3K4me3 and H3K4me3Q5dop from SU-DIPGXIII cell CUT&RUN profiling demonstrating co-localization of H3 dopaminylated chromatin with pS2-Pol2 and with published H3K27M profiles. **b-f**, Genome browser tracks showing bivalent H3K4me3/H3K27me3 chromatin profile and low H3K4me3Q5dop enrichment at the *CALCA* (**b**) and *CALCR* genes (**c**) or showing co-enrichment of H3K27M, H3K4me3, and H3K4me3Q5dop CUT&RUN signals at genes encoding Pol2 regulators (*CDK12*, *TADA1*, and *UBTF*, **d**-**f**).

#### Extended Data Figure 2

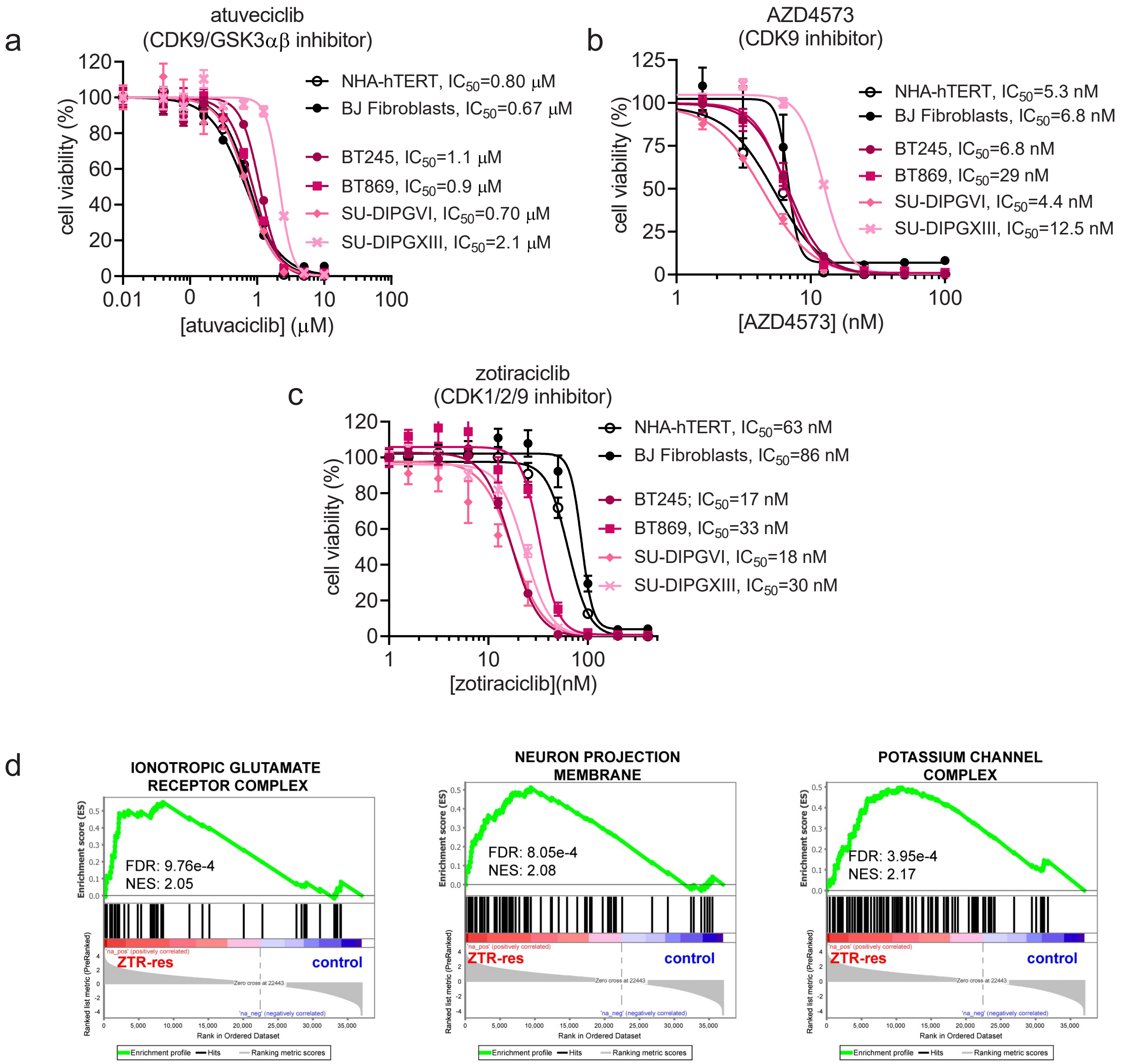

**Extended Data Fig. 2: Neuronal signaling genes are upregulated in CDK9i-resistant DMG xenograft tumors. a-c**, Normalized viability of DMG cells (magenta curves) versus non-transformed BJ fibroblasts and telomerase immortalized astrocytes (NHA-hTERT; black curves) treated with increasing doses atuveciclib (**a**), AZD4573 (**b**), or zotiraciclib (ZTR, **c**) to inhibit CDK9. Cell viability in panels **a-c** was determined using the CellTiter-Glo assay after 7 days of treatment, and error bars show +/- S.E.M. of 3-6 biological replicates. **d**, GSEA of bulk RNA-seq data from ZTR-resistant (ZTR-res) SU-DIPGXIII xenograft tumors showing increased expression of gene sets related to ion channel activity and glutamate signaling.

### Extended Data Figure 3

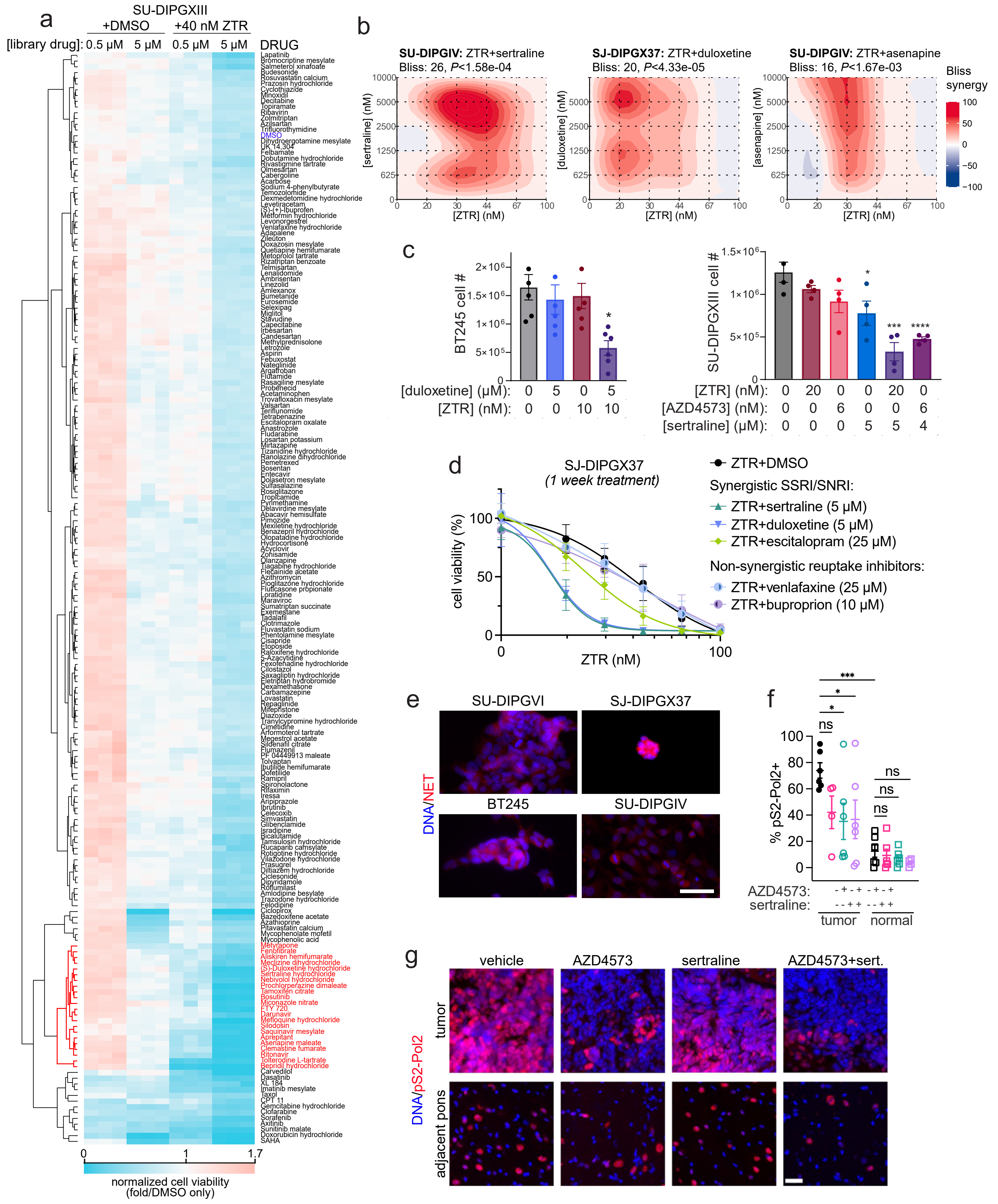

**Extended Data Fig. 3: FDA-approved drugs targeting neurotransmitter signaling sensitize DMG to CDK9i.** **a**, Heatmap of normalized SU-DIPGXIII cell viability after 7 days of treatment with 0.5 µM or 5 µM of an FDA-approved drug library combined with vehicle (DMSO; shown in blue) or 40 nM ZTR ($IC_{25}$ dose). ZTR-synergistic drug hits are highlighted in red. **b**, Heatmaps of Bliss synergy scores from SU-DIPGIV treated for 7 days with ZTR combined with either an SSRI (sertraline, left panel), an SNRI (duloxetine, middle panel) or a neurotransmitter GPCR antagonist (asenapine, left panel). **c**, Average live SU-DIPGXIII or BT245 cell counts after 7 days of treatment with ZTR (10 nM, 20 nM), AZD4573 (6 nM), sertraline (5 µM, 4 µM), and duloxetine (5 µM) alone or in combination. **d**, SJ-DIPGX37 cell viability after 7 days treatment with increasing doses of ZTR combined with vehicle (DMSO), synergistic SSRI/SNRI (sertraline, duloxetine, escitalopram), or non-synergistic neurotransmitter reuptake inhibitors (venlafaxine, bupropion). **e**, Representative images of immunofluorescence staining for the norepinephrine transporter (NET) in DMG cell lines (SU-DIPGVI, SJ-DIPGX37, BT245, SU-DIPGIV). **f,g,** Quantification (**f**) and representative images (**g**) of pS2-Pol2+ cells within the tumor region and adjacent pons of SU-DIPGXIIIP* xenograft mice treated with vehicle, AZD4573, sertraline, or AZD4573+sertraline (AZD+sert.). Scale bar is 50 µm.

#### Extended Data Figure 4

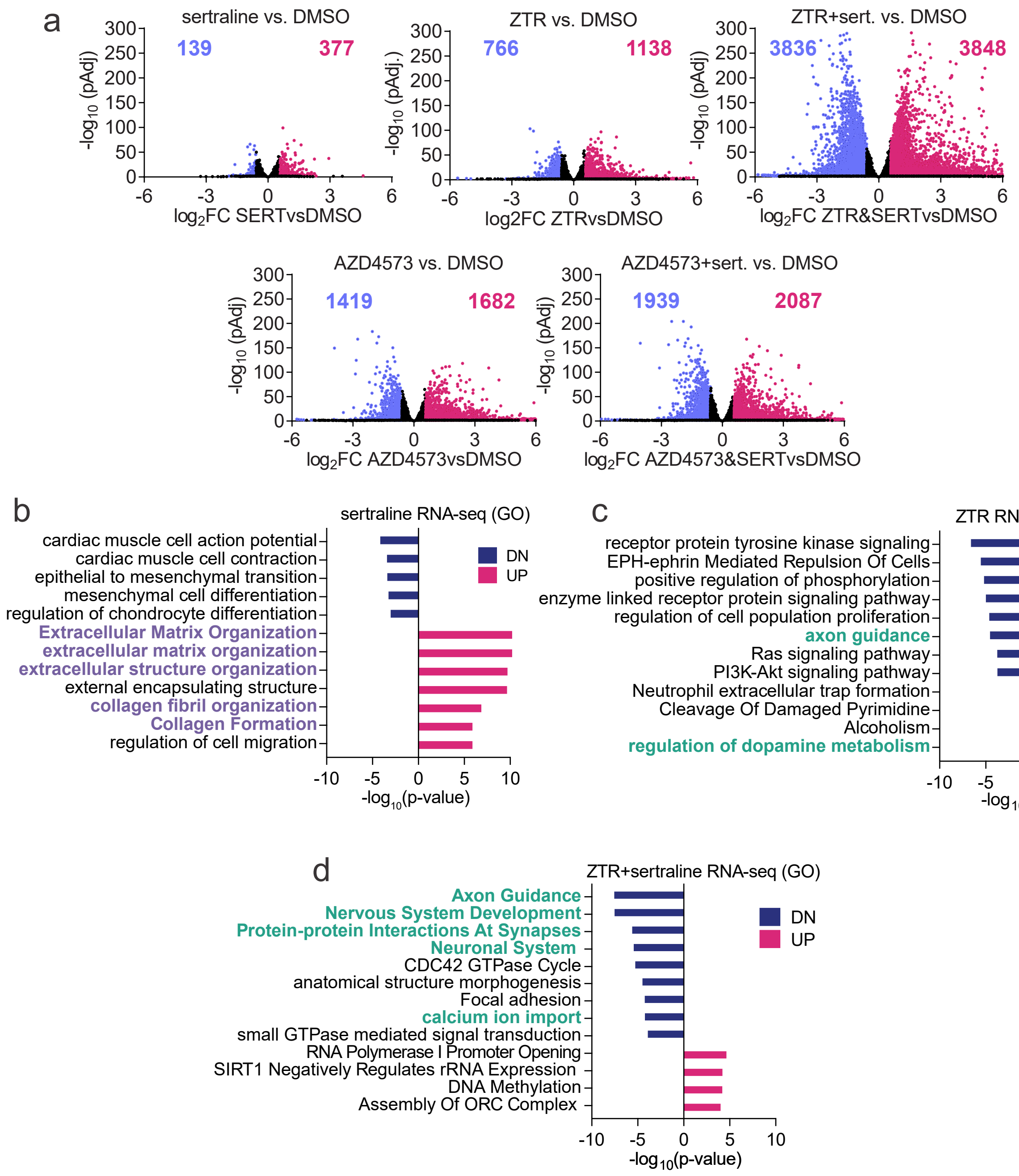

**Extended Data Fig. 4: CDK9i+SSRI co-treatment silences oncogenic and neuronal signaling genes enriched in CDK9i-resistant DMG. a**, Volcano plots summarizing RNA-seq data from SU-DIPGXIII cells treated for 24 hours with vehicle (DMSO) or combinations of 5 µM sertraline (sert.) and 20 nM ZTR , or with 6 nM AZD4573 alone or in combination. The plots display log fold change ($\log_2$FC) on the x-axis and the -$\log_{10}$(adjusted *p*-value) shown on the y-axis. **c**, Bar plots of DEG-associated GO terms in SU-DIPGXIII cells after treatment with sertraline, ZTR, or ZTR+sert.

#### Extended Data Figure 5

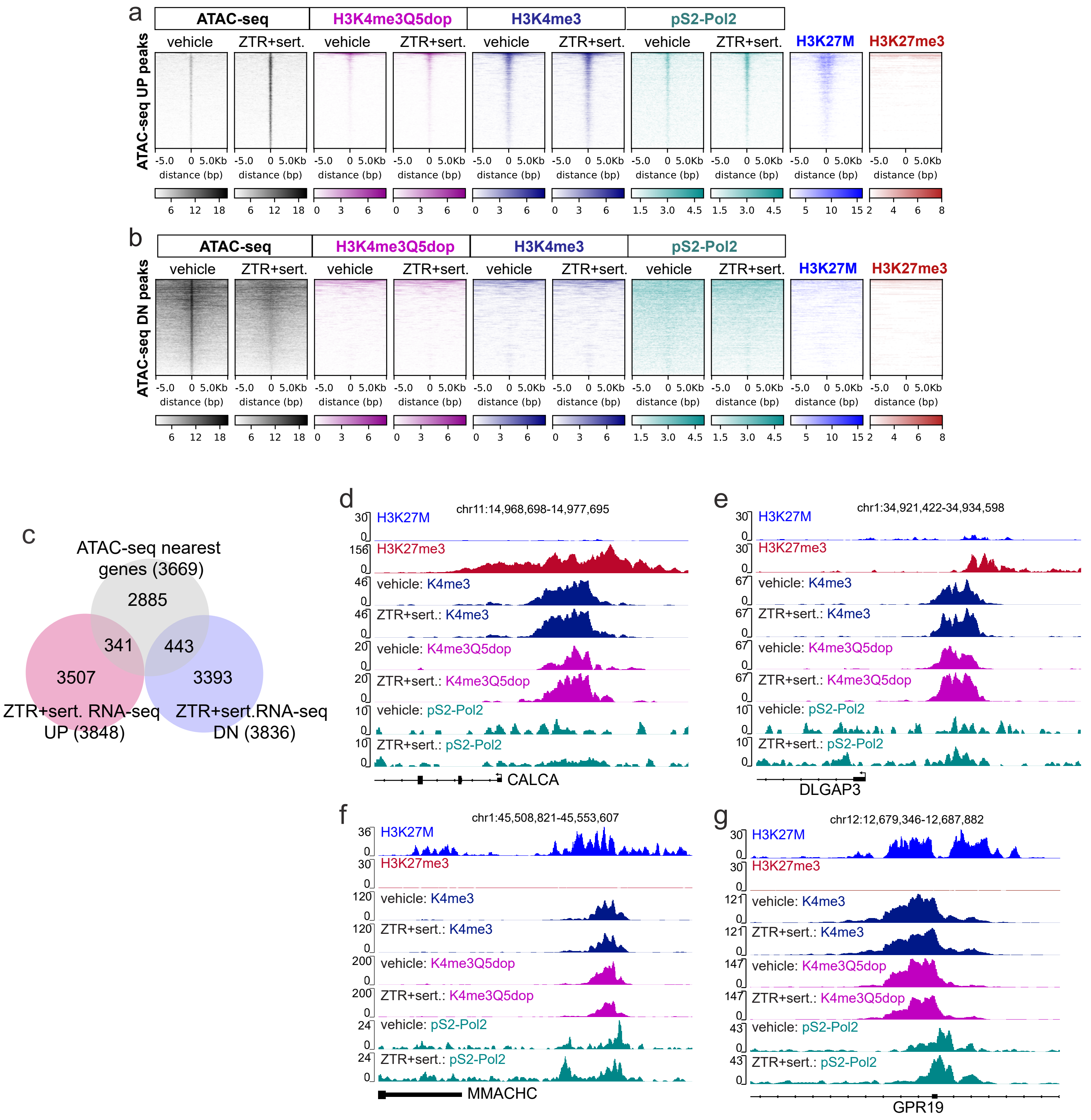

**Extended Data Fig. 5: CDK9i+SSRI treatment alters H3 dopaminylation at neuronal signaling and brain development genes. a**,**b**, Heatmaps and profile plots of increased (**a**) and decreased (**b**) ATAC-seq peaks from SU-DIPGXIII cells treated with vehicle or ZTR+sertraline showing co-localization with H3K4me3Q5dop, H3K4me3, pS2-Pol2, H3K27M, and H3K27me3. **c**, Venn diagram showing the overlap between genes nearest to differential ATAC-seq peaks ($P<0.01$) as determined using GREAT and significant differentially expressed genes (DEGs; $P<0.05$) from RNA-seq analysis of SU-DIPGXIII cells treated with ZTR+sertraline compared to DMSO control. **d**-**g**, Genome browser tracks showing ZTR+sertraline treatment-induced H3K4me3Q5dop signaling at bivalent (H3K4me3/H3K27me3) gene loci (*CALCA* and *DLGAP3*, **d**,**e**), or ZTR+sertraline treatment-repressed H3K4me3Q5dop signal at H3K27M-occupied genes (*MMACHC* and *GPR19*, **f**,**g**).

### Extended Data Figure 6

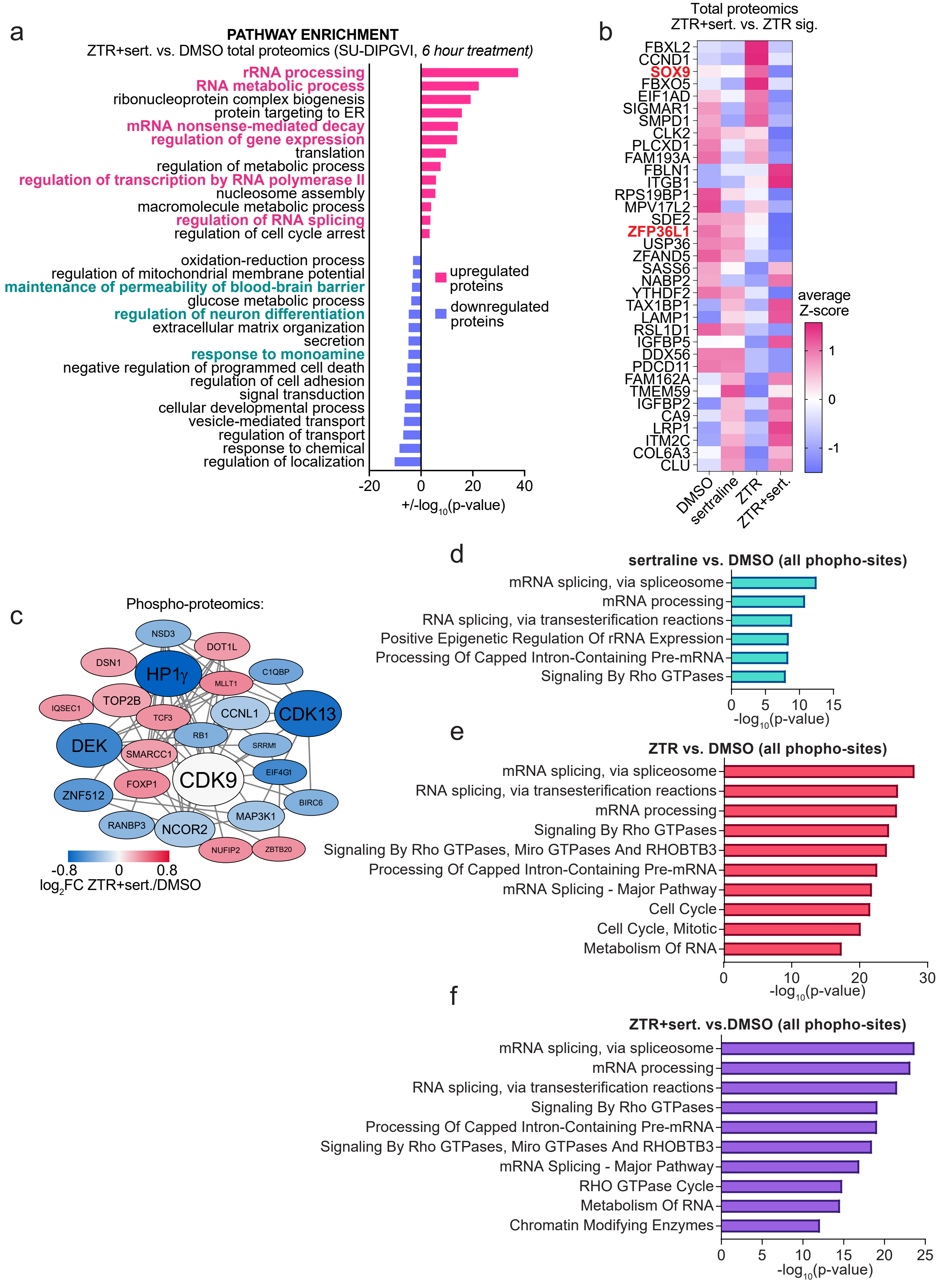

**Extended Data Fig. 6: Combining ZTR with SSRI induces distinct downstream kinase signaling events. a**, Gene ontology (GO) analysis of SU-DIPGVI total proteomics data showing reduced abundance of proteins related to the nervous system and monoamine response (green) and increased abundance of proteins involved in RNA processing and transcription (magenta) due to ZTR+sertraline (ZTR+sert.) treatment. **b**, Heatmap summarizing z-scores for top differential proteins across DMSO-, ZTR-, sertraline-, and ZTR+sert.-treated SU-DIPGVI cells. Transcription factors previously linked to glioma progression are highlighted in red. **c**, CDK9-associated protein-protein interaction network colored according to the $\log_2$ fold change ($\log_2$FC) of differential phospho-proteins in ZTR+sert.-treated compared to DMSO-treated SU-DIPGVI cells and scaled according to the adjusted p-values. **d-f**, Significant GO terms associated with all differential phospho-sites relative to DMSO control for sertraline alone (**d**), ZTR alone (**e**), or ZTR+sert. Treatment (**f**) showing $-\log_{10}$(*p*-value) for each term (**d**,**e**) or normalized enrichment ratios (**f**) for each gene category.

### Extended Data Figure 7

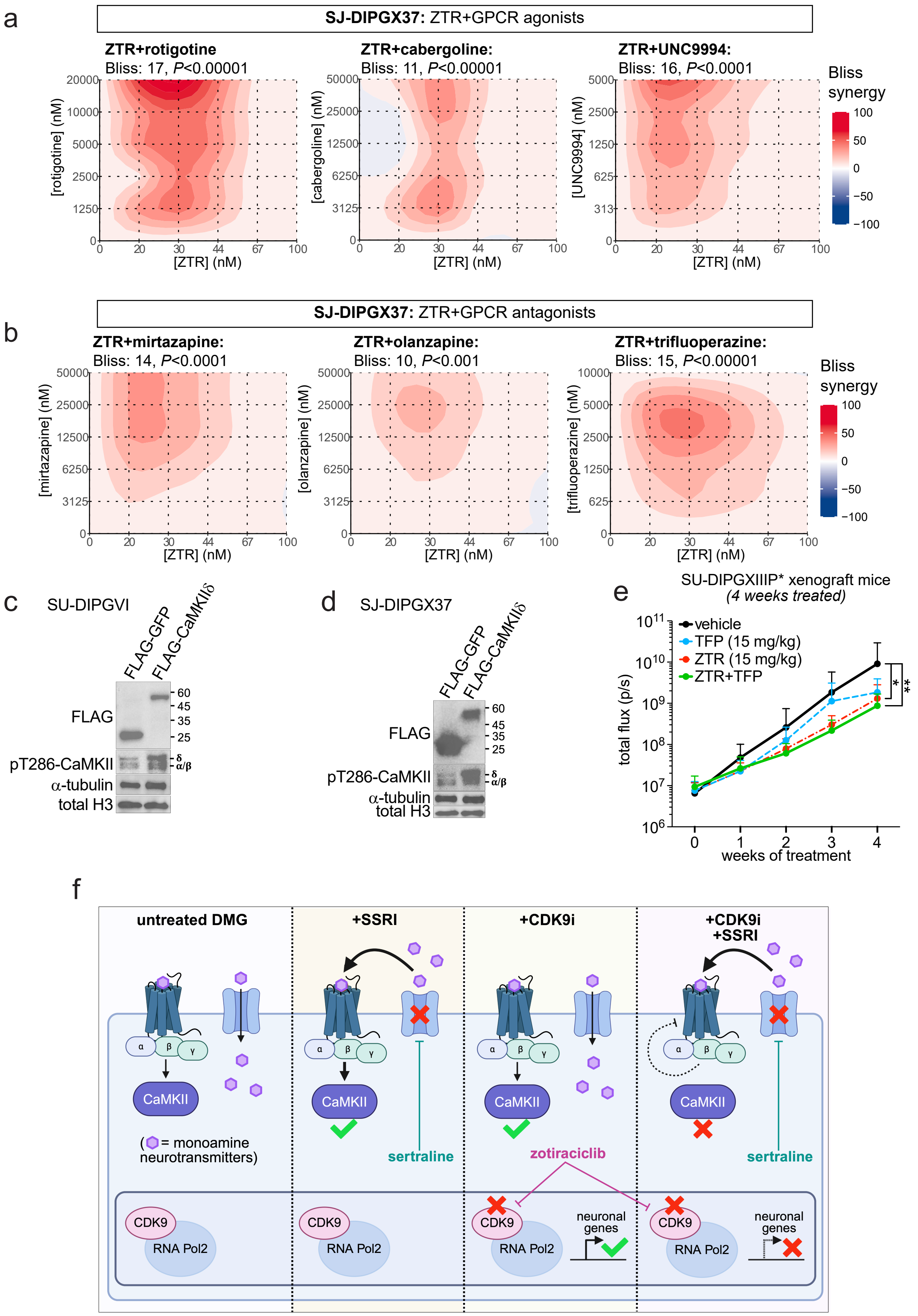

**Extended Data Fig. 7: Neurotransmitter GPCR agonists and antagonists synergize with CDK9i by reducing CaMKII activity. a,b**, Bliss synergy scores calculated from dose curve matrices of SJ-DIPGX37 cell viability data collected after 7 days of treatment with ZTR combined with neurotransmitter GPCR agonists (rotigotine, cabergoline, UNC9994, **a**) or neurotransmitter GPCR antagonists (olanzapine**,** mirtazapine, trifluoperazine, **b**). **c**,**d**, Western blot validation of flag-tagged GFP or CaMKIIδ expression in SU-DIPGVI (**d**) or SJ-DIPGX37 cells (**e**) showing increased pT286-CaMKII (a CaMKII activation marker) in FLAG-CAMK2D-transduced cells. **f**, Total tumor cell BLI flux (photons/second) from SU-DIPGXIIIP* xenograft mice treated with vehicle, TFP, ZTR, or ZTR+TFP (15 mg/kg each drug, IP) twice per week for 4 weeks. **g**, Model of ZTR+sertraline treatment-induced changes in GPCR signaling, CaMKII activation, and neuronal gene expression in DMG.

**Supplementary Table 1:** H3K4me3Q5dop and H3K4me3 CUT&RUN differential peaks in SU-DIPGXIII cells

**Supplementary Table 2:** Differentially expressed genes in ZTR-resistant xenografts and CDK9 protein correlations in patient tumors

**Supplementary Table 3:** ZTR-synergy drug screen cell viability data summary

**Supplementary Table 4:** Differentially expressed genes in SU-DIPGXIII treated with CDK9i and sertraline

**Supplementary Table 5:** pS2-Pol2 CUT&RUN, ATAC-seq, H3K4me3Q5dop, and H3K4me3 CUT&RUN differential peaks in ZTR+sert. versus vehicle-treated SU-DIPGXIII cells

**Supplementary Table 6**: Proteomics data and PhosphoSite Plus analysis in SU-DIPGVI treated with ZTR and sertraline
